# An intricate balancing act: Upstream and downstream frameshift co-regulatory elements

**DOI:** 10.1101/2024.06.27.599960

**Authors:** Samuel Lee, Shuting Yan, Abhishek Dey, Alain Laederach, Tamar Schlick

**Affiliations:** Department of Chemistry, New York University, New York, 10003, NY, U.S.A; Department of Biotechnology, National Institute of Pharmaceutical Education and Research-Raebareli (NIPER-R), Lucknow, 226002, Uttar Pradesh, India; Department of Biology, University of North Carolina at Chapel Hill, Chapel Hill, 27599, NC, U.S.A; Courant Institute of Mathematical Sciences, New York University, New York, 10012, NY, U.S.A; NYU-ECNU Center for Computational Chemistry, NYU Shanghai, Shanghai, 200062, P.R.China; NYU Simons Center for Computational Physical Chemistry, New York University, New York, 10003, NY, U.S.A

**Keywords:** RNA, frameshift, graph theory

## Abstract

Targeting ribosomal frameshifting has emerged as a potential therapeutic intervention strategy against Covid-19. During ribosomal translation, a fraction of elongating ribosomes slips by one base in the 5^′^ direction and enters a new reading frame for viral protein synthesis. Any interference with this process profoundly affects viral replication and propagation. For Covid-19, two RNA sites associated with ribosomal frameshifting for SARS-CoV-2 are positioned on the 5^′^ and 3^′^ of the frameshifting residues. Although much attention has been on the 3^′^ frameshift element (FSE), the 5^′^ stem-loop (attenuator hairpin, AH) can play a role. The formation of AH has been suggested to occur as refolding of the 3^′^ RNA structure is triggered by ribosomal unwinding. However, the attenuation activity and the relationship between the two regions are unknown. To gain more insight into these two related viral RNAs and to further enrich our understanding of ribosomal frameshifting for SARS-CoV-2, we explore the RNA folding of both 5^′^ and 3^′^ regions associated with frameshifting. Using our graph-theory-based modeling tools to represent RNA secondary structures, “RAG” (RNA-As-Graphs), and conformational landscapes to analyze length-dependent conformational distributions, we show that AH coexists with the 3-stem pseudoknot of the 3^′^ FSE (graph 3_6 in our dual graph notation) and alternative pseudoknot (graph 3_3) but less likely with other 3^′^ FSE alternative folds (such as 3-way junction 3_5). This is because an alternative length-dependent Stem 1 (AS1) can disrupt the FSE pseudoknots and trigger other folds. In addition, we design four mutants for long lengths that stabilize or disrupt AH, AS1 or FSE pseudoknot to illustrate the deduced AH/AS1 roles and favor the 3_5, 3_6 or stem-loop. These mutants further show how a strengthened pseudoknot can result from a weakened AS1, while a dominant stem-loop occurs with a strengthened AS1. These structural and mutational insights into both ends of the FSE in SARS-CoV-2 advance our understanding of the SARS-CoV-2 frameshifting mechanism by suggesting a sequence of length-dependent folds, which in turn define potential therapeutic intervention techniques involving both elements. Our work also highlights the complexity of viral landscapes with length-dependent folds, and challenges in analyzing these multiple conformations.

## Introduction

The COVID-19 pandemic and its lasting effects have inspired innovative research into coronavirus proteins and viral RNAs, as well as underscored other potential viral threats. The frameshifting process has been a promising focus area of research with long-term therapeutic potential against viral infections (1; 2; 3; 4; 5; 6). RNA frameshifting is regulated by the FSE of the RNA virus. This element is located in the open reading frame ORF1a,b region of SARS-CoV-2’s viral genome that codes for the polyproteins necessary for viral protein synthesis (Figure 1). However, ORF1a and ORF1b overlap by a single nucleotide in coronavirus genomes, with ORF1b starting from the −1 reading frame compared to ORF1a. The FSE is responsible for the Programmed −1 Ribosomal Frameshift (−1 PRF), where translating ribosomes shift their reading frames by one nucleotide in the 5^′^ direction (−1). After this frameshift is complete, the ribosomes can decode the ORF1b polyproteins (7).

**Fig. 1.**
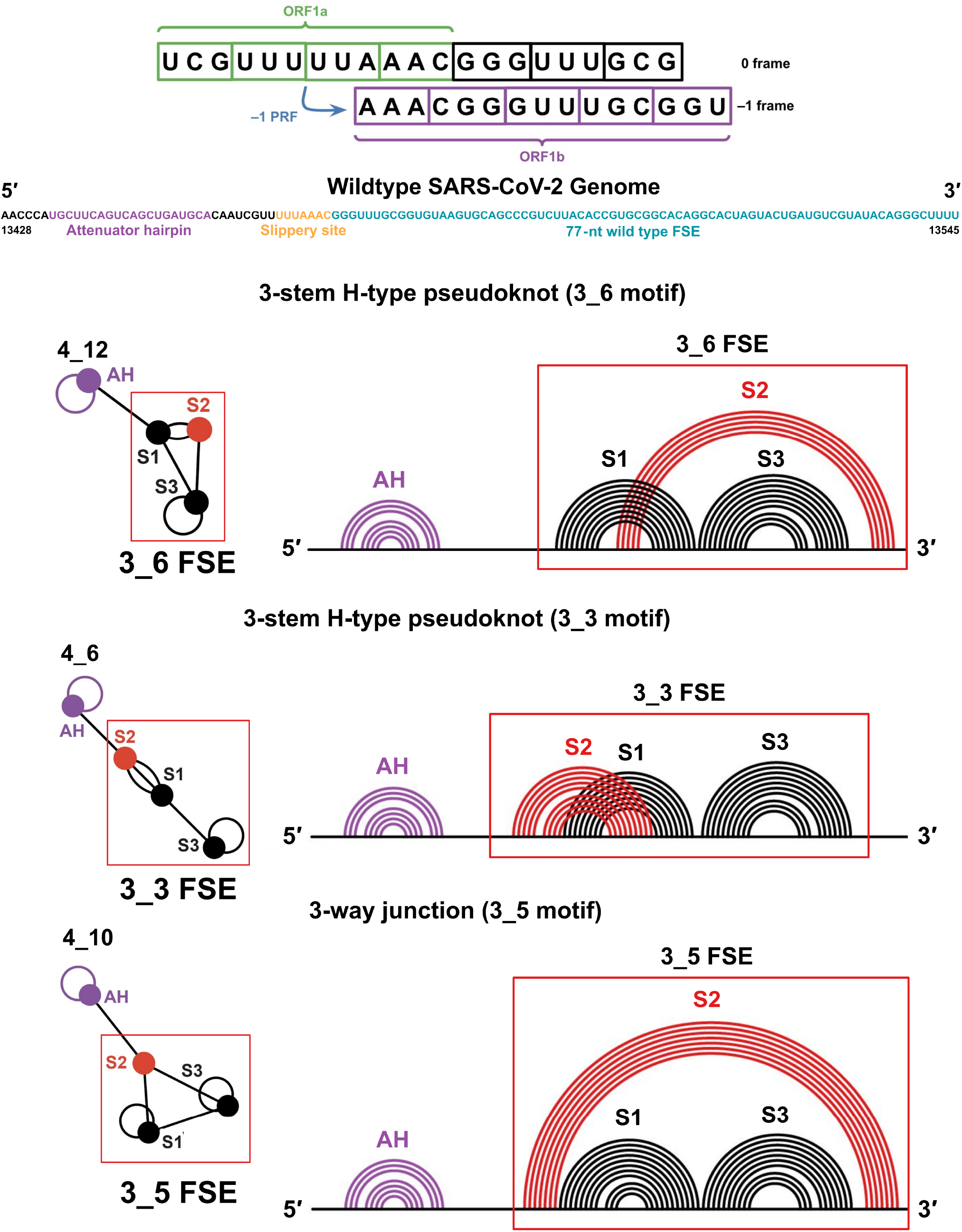
SARS-CoV-2 FSE sequence, and three associated 2D motifs of 118-nt FSE-containing RNA folds emerging from this work. To illustrate the frameshifting process, the 0-frame and −1-frame codons are labeled with the overlapping regions of ORF1a and ORF1b marked. The attenuator hairpin, the 7-nt slippery site and downstream stimulatory pseudoknot are highlighted. For each 2D structure containing AH and FSE that adopts one of three possible folds, including 3-stem H-type 3_6 pseudoknot, alternative 3_3 pseudoknot, and three-way junction 3_5 (unknotted RNA), corresponding dual graphs and arc plots are shown, with color coded stems and loops labeled.

The FSE of coronavirus has been characterized as a pseudoknot and associated with the −1 PRF slippage event at the 7-residue slippery site (8; 9; 10; 11; 12; 13). The SARS-CoV-2 FSE is a 84 nucleotides segment and contains, from 5^′^ to 3^′^, a 7-nt slippery site (UUUAAAC) and a 77-nt downstream RNA structure (see Figure 1).

Many different models suggest how the pseudoknot acts in translation. The “torsional restraint” model suggests that the ribosome unwinding Stem 1 of the pseudoknot (moving in the forward direction) induces stress by supercoiling Stem 2. This resistance is enough to counteract the ribosome’s forward movement, causing the ribosome’s A- and P-sites to essentially pause at the slippery site, setting the ribosome up for the frameshift (12). The “9 Å Solution” model suggests that the downstream RNA impedes ribosomal translation and stretches the 8-residue spacer between the slippery site and the pseudoknot.

A 20-residue attenuator hairpin (AH) upstream of the FSE may be involved in frameshifting as well (14; 15). The AH may downregulate the frameshifting process, causing ribosomes to dissociate from the mRNA before receiving the −1 PRF signal. Thus, the AH controls the ratio of ORF1a to ORF1b polyprotein products, thereby helping optimize viral protein synthesis and replication (16; 17).

Much remains unclear about how the FSE and AH regions work in tandem. Here, we aim to unravel how AH and FSE regions fold together and influence frameshifting.

In our previous work, we explored the multiple-conformational landscape of the SARS-CoV-2 downstream FSE using our graph-theoretic approach, “RAG” (RNA-As-Graphs), which uses tree and dual graphs to represent, study, predict, and design RNA secondary (2D) structures (18; 19; 20; 21; 22). With our coarse-grained dual graphs, we delineated the alternative RNA structures of the downstream FSE and designed mutants via inverse folding (23) that transform the FSE topology into others (24; 25). Our FSE landscape includes the dominant 3-stem pseudoknot, which we term dual graph 3_6 (confirmed by X-ray, NMR and Cryo-EM (26; 27; 28; 2; 29)), the alternative pseudoknot 3_3, and the three-way junction 3_5 (boxes in Figure 1) (25).

The three RNA motifs in Figure 1 contain the same Stems 1 and 3, but different Stem 2, which involves the 3^′^-end (in 3_6) or 5^′^-end (in 3_3). The 3_6 pseudoknot has also been confirmed by various methods (26; 27; 28; 2; 29) mostly for downstream sequences. Unknotted structures have been identified (30; 31; 32; 33; 34), which coexist with AH in long sequence constructs. We have proposed that structural transitions among these three (and other possible) motifs likely exist and play an important role in frameshifting (35). In our first FSE work (24), we designed four double mutants for 77-nt RNAs that transform the 3_6 pseudoknot into stem-loops 2_1 and 3_2, the three-way junction 3_5 and a 3_3 pseudoknot with S2 and S3 intertwined (rather than S1 and S2 in 3_3 in Figure 1). These 77-nt mutants were embedded in 114-nt RNAs called M1-M4 in Ref (36) and experimentally tested by DMS-MaP. Later in Ref (25) we also designed 77-nt minimal mutants (2 to 6 residues changed) to strengthen the 3_3, 3_6 and 3_5 FSE structures (termed M3_3, M3_6, M3_5, respectively (25), see mutant sequences in Table 1).

**Table 1.**
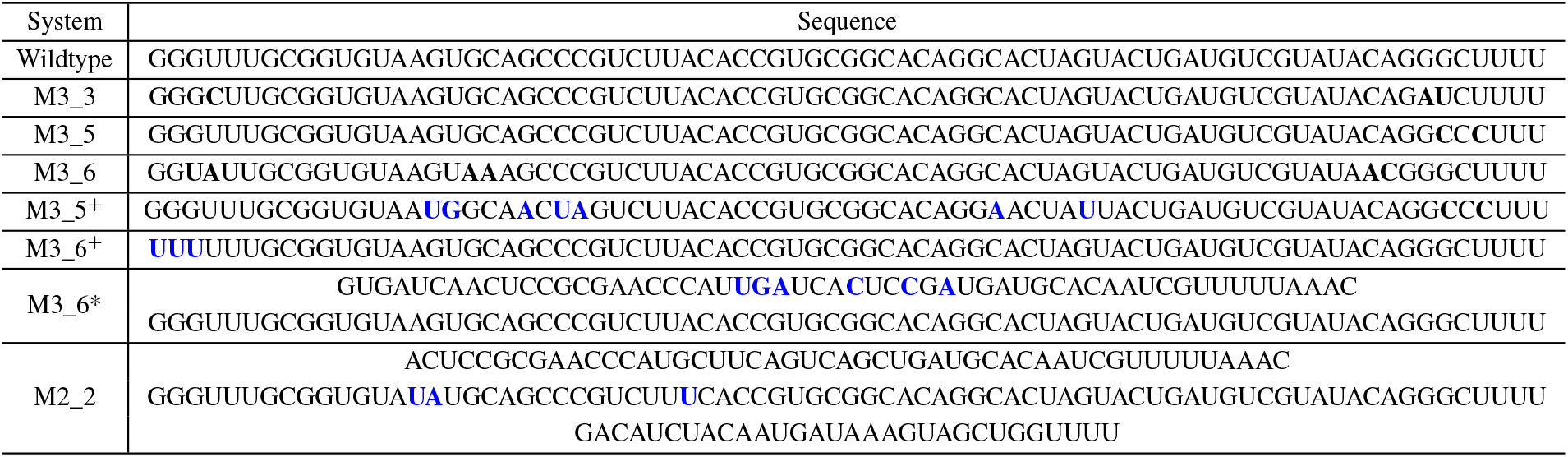
Sequences of central 77-nt FSE and long FSE-containing constructs in the mutants studied in this work. Mutations from Ref (25) (M3_3, M3_6 and M3_5) are in bold and red. Mutations designed in this work (M3_5^+^, M3_6^+^, M3_6* and M2_2) are in bold and blue.

DMS-MaP data in Ref (36) for 114-nt versions of our four earlier 77-nt mutants (24) yield 2_1, 3_2 and 3_5 as predicted for M1–M3, but 2_1 for the FSE-M4 mutant instead of the 3_3 pseudoknot due to a flexible Stem 3 loop that cannot be engaged in a pseudoknot (see Results in Ref (36) and Figure S1). Pekarek et al. (36) also reported that our earlier mutants decreased frameshifting by an order of magnitude (24) (including M3_5 mentioned here) (from 25.6% to 1∼1.3%). More recent experiments suggest that our structure-stabilizing and transition-suppressing mutants reduce frameshifting significantly (37), confirming our hypothesis that structural changes in FSE stem formation and overall folding play a significant role in fine-tuning the frameshift process.

Here we continue to explore the length-dependent RNA folding via graph theory-based modeling to discover the relationship between AH formation and various alternative forms of the downstream stimulatory sequence. Are the three different 77-nt FSE motifs (3_6, 3_3, and 3_5) stable when the upstream AH forms at longer RNA lengths? To answer this, we predict RNA folds as sequence increases, thereby defining *conformational landscapes*, which we introduced in (25; 35), and as shown in Figure 2.

**Fig. 2.**
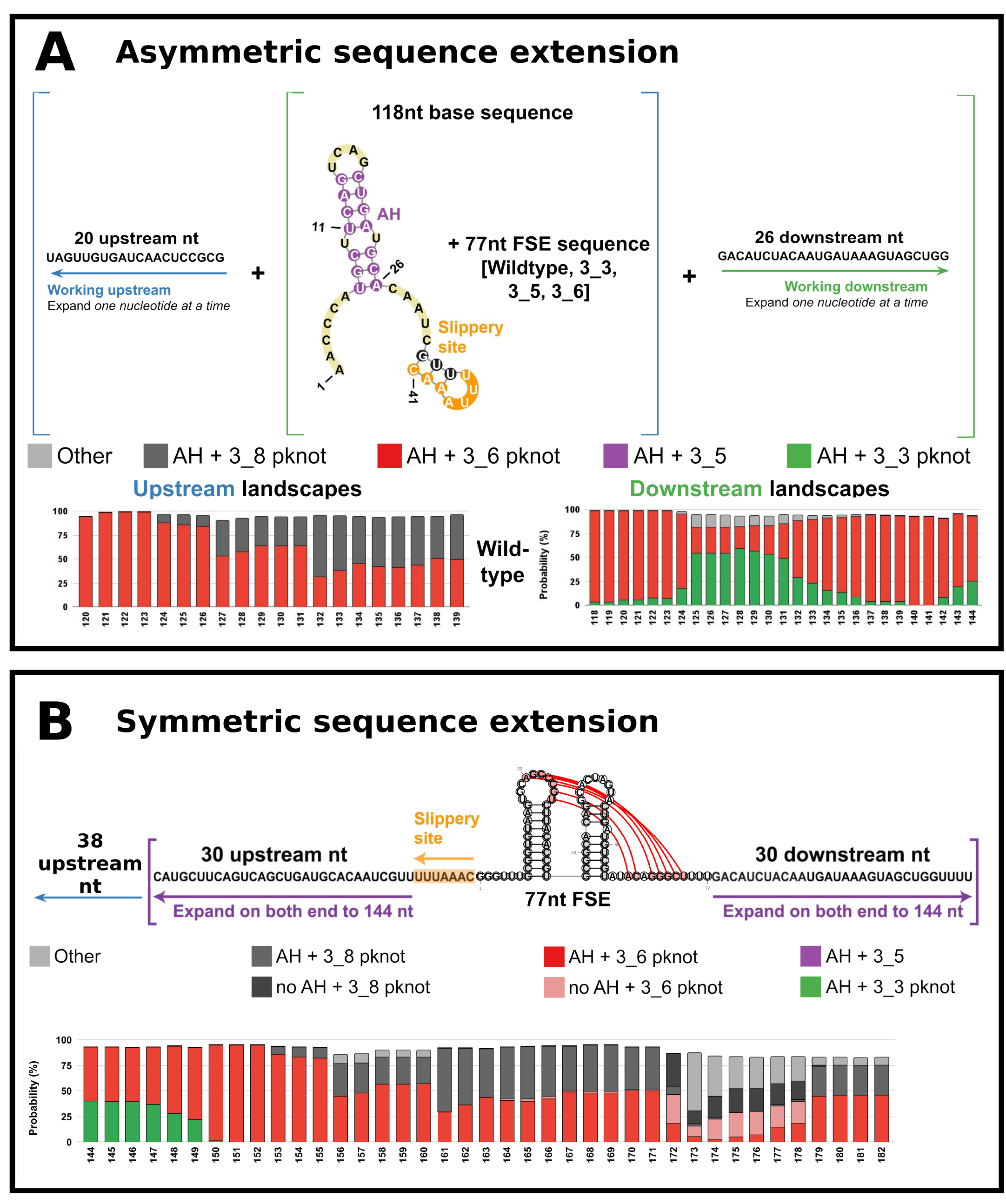
Conformational landscapes of the wildtype FSE with upstream or downstream extensions of sequence (top) and symmetric extension of sequence (bottom). (A) In the conformational landscapes with increasing upstream or downstream sequence, the ensemble of 2D folds at each sequence length is predicted by NUPACK (41). For each length, probabilities of 2D folds are calculated from the Boltzmann factor, and all structures containing AH and 3_6, 3_3 or 3_5 motifs are individually summed. Motifs containing 3_6 and AH are in red; motifs containing 3_3 and AH are in green; and motifs containing 3_5 and AH are in purple. See Figure 3 for examples of folds at different lengths. (B) The bottom landscape is computed with symmetric extension on both ends of the 144-nt segment containing the FSE.

We find that the 3_3 or 3_6 pseudoknot FSE topology can coexist with AH in the wildtype sequence, while the 3_5 motif is not likely compatible at long sequences. The pseudoknots 3_3 and 3_6 of (downstream) FSE likely form when the ribosome unwinds AH. The switch between 3_6 and 3_3 depends on whether the residues upstream or downstream the 77-nt FSE are exposed. In long sequence constructs, upstream and downstream sequences form long and stable anchoring stems, such as AS1, thereby blocking pseudoknot formation in the downstream FSE. The 77-nt 3_3 and 3_6 pseudoknots strengthened by our M3_3 and M3_6 mutations are stable when AH exists. However, a stable three-way junction 3_5 is not compatible with AH, and requires additional mutations to retain 3_5 with an upstream AH. We also design two new mutants, to either block AH formation or the pseudoknot, supporting the mechanism obtained from our landscapes.

Together, our conformational landscapes, mutant designs, and new published experimental data help describe the structural relation between upstream AH and downstream FSE, and outline a mechanistic picture of ribosomal frameshifting involving multiple ribosomes and a dynamic folding/unfolding cascade during translation. Key in our mechanism and associated frameshifting efficiency are structural switches between unknotted stem-loops (when AS1 forms) and 3-stem pseudoknot/junction during the folding/refolding cycles. Thus, our motif strengthening mutants which suppress refolding transitions define new avenues for targeting ribosomal frameshifting.

## Materials and Methods

### RAG dual graphs and inverse folding

To represent RNA 2D structures in our RNA-As-Graphs framework, double-stranded regions (stems) are denoted vertices, and single-stranded regions (bulges, loops and junctions) are edges in dual graphs (18; 38; 39). Hairpin loops are represented as self edges, and unpaired residues at the 5^′^ and 3^′^ ends are ignored.

We use our inverse-folding protocol RAG-IF (23) to mutate the M3_5 FSE sequence to fold onto dual-graph motifs containing the AH and a stable 3_5 in the downstream stimulatory FSE region. RAG-IF has three steps: 1) identify mutation regions and target 2D structures, 2) produce candidate sequences by mutations using a genetic algorithm (GA), and 3) optimize the mutated sequence pool by sorting and retaining only minimal or essential mutations that fold onto the target graph. GAs mimic evolution in nature, where the template sequence undergoes iterations of random mutation, crossover, and selection, and those with high fitness are retained. The fitness is determined by the Hamming distance. The nominated sequences are folded by IPknots (40), and further examined by NUPACK (41). Sequences that achieve the target folding are further optimized to obtain the minimal mutations by removing the non-essential mutations (23).

### Selection of mutation regions

We compare the wildtype FSE graph and the target graph to identify the smallest possible mutation regions required for the transformation. Considering that most of the motifs generated from appending downstream residues to the M3_5 mutant contain a 3_3 FSE, we focused on breaking the 3_3 pseudoknot to allow the 3_5 FSE to form in its place. We specifically selected the downstream residues of 3_3 S2 as the mutation site. Here, we describe how this is accomplished for our eight most successful target graphs: 6_458, 5_104, 5_90, 6_481, 7_2207, 6_390, 4_6, and 7_2365. We focus on the motif topology and allow the lengths of stems and loops to vary.

### Conformational landscape calculation

For each variant (wildtype, PSM M3_3, M3_5, M3_6), we start with the base 41-nt sequence: AACCCAUGCUUCAGUCAGCUGAUGCACAAUCGUU-<u>UUUAAAC</u>.

The base 41-nt sequence contains the AH and the slippery site. The sequence is selected according to Pekarek et al. (36) FSE-V2 construct. We append the 77-nt unique sequence corresponding to each variant (see Table 1) after the slippery site (UUUAAAC, see underlined above), see full sequence of 118-nt in Table S1. Working downstream, we add the 26-nt sequence GACAUCUACAAUGAUAAAGUAGCUGG one nucleotide at a time, appending it to the 3^′^-end of each template sequence (base 41-nt + unique 77-nt). Working upstream, we add the 20-nt sequence UAGUUGUGAUCAACUCCGCG one nucleotide at a time, appending it to the 5^′^-end of each template sequence.

For each coronavirus FSE, we use NUPACK v3.2.2 (41; 42) *subopt* mode and option *-pseudo* to predict RNA secondary structure ensembles for lengths from 118 to 138-nt adding upstream sequence and lengths from 118 to 144-nt adding downstream sequence. The output contains the predicted 2D structures together with their free energy estimates. We then use the Boltzmann distribution to compute the partition function *Z* and the probability *p*_*i*_ for each 2D structure *i*,

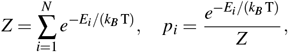

where *E*_*i*_ is the free energy estismate of structure *i, k*_*B*_ is the Boltzmann constant, and T is the room temperature (37 °C).

For each length, we apply our dual graph representation and sum up probabilities of structures that correspond to the same graph. Only graphs with probabilities ≥ 1% are retained, and representative 2D structures are recorded for these graphs.

Next, we identify the minimal motifs within these example structures that involve the 118-nt FSE-containing regions. Once we identify all minimal motifs, we sum up probabilities of dual graphs that correspond to the same minimal motif, and retain only motifs with probabilities ≥ 5%.

### DMS-MaP chemical probing

#### In vitro RNA chemical probing read by mutational profiling

156-nt SARS-CoV-2 FSE-containing construct was synthesized as G-blocks from Integrated DNA Technologies (IDT). The whole construct was flanked at both 5^′^ and 3^′^-end by RNA cassettes (43). For *in vitro* transcription of the 156-nt FSE-containing construct, a T7 promoter region was added to its 5^′^-end. Transcription was performed using T7 HiScribe RNA synthesis kit (New England Biolabs). Generated RNA was subjected to DNase treatment (TURBODNase) which was further purified using Purelink RNA mini kit (Invitrogen) and quantified using nanodrop. Standardized cassettes used are specifically designed to form hairpin structures upstream and downstream of the construct, which have been shown to allow more accurate structure predictions based on the DMS data as the amplicon primers do not cover the signal for the 5^′^ and 3^′^ ends of the construct.

For chemical probing, 6 *µ*g of purified synthetic RNA was denatured at 65 °C for 5 min and snap-cooled in ice. Following denaturation, folding buffer (100 mM KCl, 10 mM MgCl2, 100 mM Bicine, pH 8.3) was added to the denatured RNA and the whole reaction was incubated at 37 °C for 10 min. The folded RNA was further treated with 10 *µ*L of 1:10 ethanol diluted Dimethyl Sulfate (DMS). For control, an equivalent volume of ethanol was added to the folded RNA. Probing was initiated at 37 °C for 5 min and was quenched afterwards using 100 *µ*L of 20% *β* -mercaptoethanol (*β* -ME). Modified and un-modified RNAs were purified using Purelink RNA mini kit (Invitrogen) and quantified using nanodrop.

#### Library construction, sequencing, and data processing

Both chemically modified and unmodified RNAs were reverse transcribed using Gene-specific primer (Table 2) complementary to 3^′^ RNA cassette and Superscript II reverse transcriptase under error prone conditions as previously described (44). cDNA generated was further purified using G50 column (GE healthcare) and subjected to second strand synthesis (NEBNext Second Strand Synthesis Module). For constructing next-generation sequencing libraries, the double-stranded (ds) cDNA was PCR amplified using the primers directed against 5^′^ and 3^′^ RNA cassettes and NEB Q5 HotStart polymerase (NEB). To introduce unique barcodes, secondary PCR was performed using TrueSeq primers (NEB) (44). Resultant libraries were purified using Ampure XP (Beckman Coulter) beads and quantified using Qubit dsDNA HS Assay kit (ThermoFisher). For quality check, libraries were subjected to Agilent Bioanalyzer 2100. Final libraries were sequenced as 2 x 151 paired end read on Illumina MiSeq platform. To calculate mutation frequency in both chemically modified (DMS treated) and control (ethanol treated) RNA samples shapemapper2 algorithm was used (45). Chemical modifications on each RNA nucleotide was calculated using the following equation:

**Table 2.**
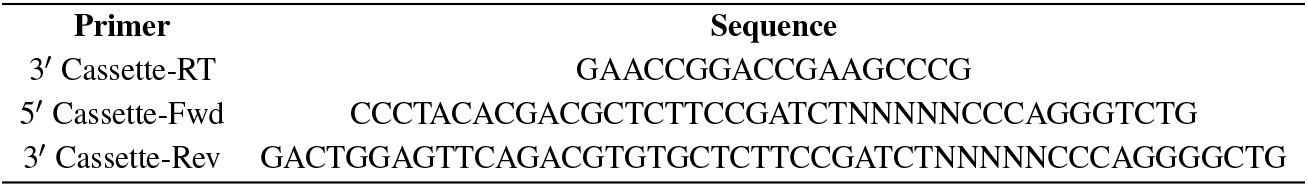
Primers used for PCR and Reverse transcription of 156-nt SARS-CoV-2 FSE-containing sequence.

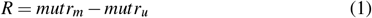

where R is the chemical reactivity, *mutr*_*m*_ is the mutation rate calculated for chemically modified RNA and *mutr*_*u*_ is the mutation rate calculated for unmodified control RNA samples (44).

### RNA structure predictions with DMS-MaPseq data

#### DREEM

Using the “detection of RNA folding ensembles using expectation–maximization” (DREEM) algorithm (46), alternative structures were directly identified from the sequencing reads from DMS-MaPseq.

Using an expectation-maximization technique, DREEM clusters the reads into discrete groups based on patterns of DMS-induced mutations. Loglikelihood is maximized to obtain the DMS modification rate per base for each cluster. In this work, a maximum of K=3 clusters were used to group the bit-vectors. The resulting DMS reactivities for each cluster were then used as constraints for ShapeKnots (47) predictions. Hence, distinct structural clusters with their relative ratios result in different folds, which represents the heterogeneity of RNA secondary structure.

#### DRACO

We also applied the DRACO algorithm (48), which performs deconvolution of alternative RNA conformations from mutational profiling experiments with a combination of spectral clustering and fuzzy clustering, to validate the structure prediction. Spectral clustering is performed for the sliding windows along the transcript, allowing the optimal number of coexisting conformations (clusters) to be automatically identified from the eigengaps. Following the determination of the number of clusters, fuzzy clustering is carried out to allow bases to be weighted according to their affinity to each cluster. DRACO then reconstructs overall mutational profiles by merging overlapping windows with the same number of clusters. DRACO reports consecutive sets of windows with varying amounts of clusters separately.

The pair-end reads were merged by pear (49) and mapped to the reference sequence using the rf-map tool (50) (parameters: -b2 -cqo -ctn - mp ‘–very-sensitive-local’). Resulting BAM files were then analyzed with the rf-count tool to produce MM files (-r -m -mm -na -ni). MM files were analyzed with DRACO (48) (parameters: –allNonInformativeToOne –nonInformativeToSurround –minClusterFraction 0.1) and deconvoluted mutation profiles were extracted from the resulting JSON files. Normalized reactivity profiles were obtained by first calculating the raw reactivity scores via the scheme by Zubradt et al. (51) as the per-base ratio of the mutation count and the read coverage at each position, and then by 90% Winsorizing as normalization method, using the rf-norm tool (50) (parameters: -sm 4 -nm 2 -rb AC -mm 1). Data-driven RNA structure prediction was performed using ShapeKnots (47) and the normalized reactivity profiles.

## Results

### Overview

In the subsequent sections, we generate conformational landscapes of FSE-containing RNA sequences computationally (see note below) to analyze the various conformations of the SARS-CoV-2 variants (wildtype, and 3_3, 3_6 and 3_5 mutants denoted as M3_3, M3_6 and M3_5) as a function of sequence length, mimicking ribosomal translation.

That is, these landscapes mimic RNA folding at increasing lengths (35). We begin with a sequence that contains AH and 3_6 FSE (also modeled by Pekarek et al. (36)). Adding *upstream* residues mimics the RNA sequence that the ribosome encounters when the genome is unwound as the ribosome moves along the RNA transcript. For comparison, we also construct landscapes with varying *downstream* sequence lengths to mimic the refolding when the ribosome moves further downstream. For added perspective, we also consider symmetric increases of sequence length, as in our previous study (25). Although our computed landscapes only suggest general trends, our prior experimental confirmation with SHAPE (25) and current comparison with DMS-MaPseq chemical probing data lends confidence in this approach.

Starting from the 118-nt base sequence, which contains the upstream AH and downstream FSE, we calculate conformational landscapes over various sequence lengths using graph IDs, which allow us to describe the conformational landscape succinctly, e.g., 20% 3_3 and 80% 3_6. Because sequences are longer than 77-nt, higher-order folds result (i.e., with more than 3 vertices), but these large graphs may contain the 3_6, 3_3 and 3_5 as subgraphs and we color them accordingly (see Figure 1). The symmetric landscapes reflect both upstream and downstream extensions.

Clearly, sequence length and context impact the folding of the AH-FSE region. For instance, increasing the length of the wildtype sequence upstream aids in the emergence of AS1, which replaces S1 and modifies the FSE topology. Expanding downstream triggers an FSE conformational transition between the two pseudoknots, 3_6 and 3_3, in the absence of an upstream region for AS1. In the symmetric expansion, we also capture transitions between the pseudoknots when downstream residues are added and competition between the 3_6 pseudoknot and other folds as in our upstream landscape.

Additionally we find that our previously designed M3_3 and M3_6 mutants (25) maintain the 3_3 and 3_6 FSEs effectively in the presence of added residues for RNAs in the 118∼138-nt range, while the 3_5 FSE easily breaks due to the formation of other structures (i.e., AS1). We apply these insights to design additional mutants: (1) a long RNA mutant (M3_5^+^) that stabilizes 3_5 using our genetic algorithm RAG-IF (see **Materials and Methods**); (2) mutants (M3_6^+^ and M3_6*) that block AS1 and AH, respectively, and maintain the 3_6 pseudoknot for long RNAs; and (3) mutant M2_2 that supports the 2_2 stem-loop for the FSE with a stable AS1 (12-bp) (see Table S1 for full sequences of the mutants studied in this work).

Here in the main text, our landscapes are generated by NUPACK (41) (see Methods), which has yielded consistent results for our work and in comparison to experimental reference before (25). NUPACK generates suboptimal structures that allow us to determine the conformational distribution (see information in SI on the performance of other packages).

In addition to our computational landscapes, our paper presents and analyzes new DMS-MaPseq FSE chemical probing data (see Methods) for 156-nt FSE-containing sequence. Previously published data are also analyzed with the mechanistic information gleaned from our landscapes.

### Wildtype SARS-CoV-2 Conformational Landscapes

We begin with a secondary-structure folding analysis as a function of sequence length on the wildtype SARS-CoV-2 sequence (see Materials and Methods). In brief, at each RNA length, we fold the RNA “in silico” and report the percentages of each “fold” (denoted by its RAG graph ID) within the conformational ensemble. The color codes in our landscapes represent our four fold classes. Graphs that include AH with 3_6, 3_3 or 3_5 as subgraphs (see Figure 3), or other folds, are colored in different colors (red, green, purple or gray). For example, starting at 118-nt, we have 95.2% of the 3_6 pseudoknot and 3.6% of the 3_3 pseudoknot (Figure 2A). Thus, the majority of the histogram bar is red (3_6 family) and the rest is green (for the 3_3 family) for this RNA length (see Figure 2A).

**Fig. 3.**
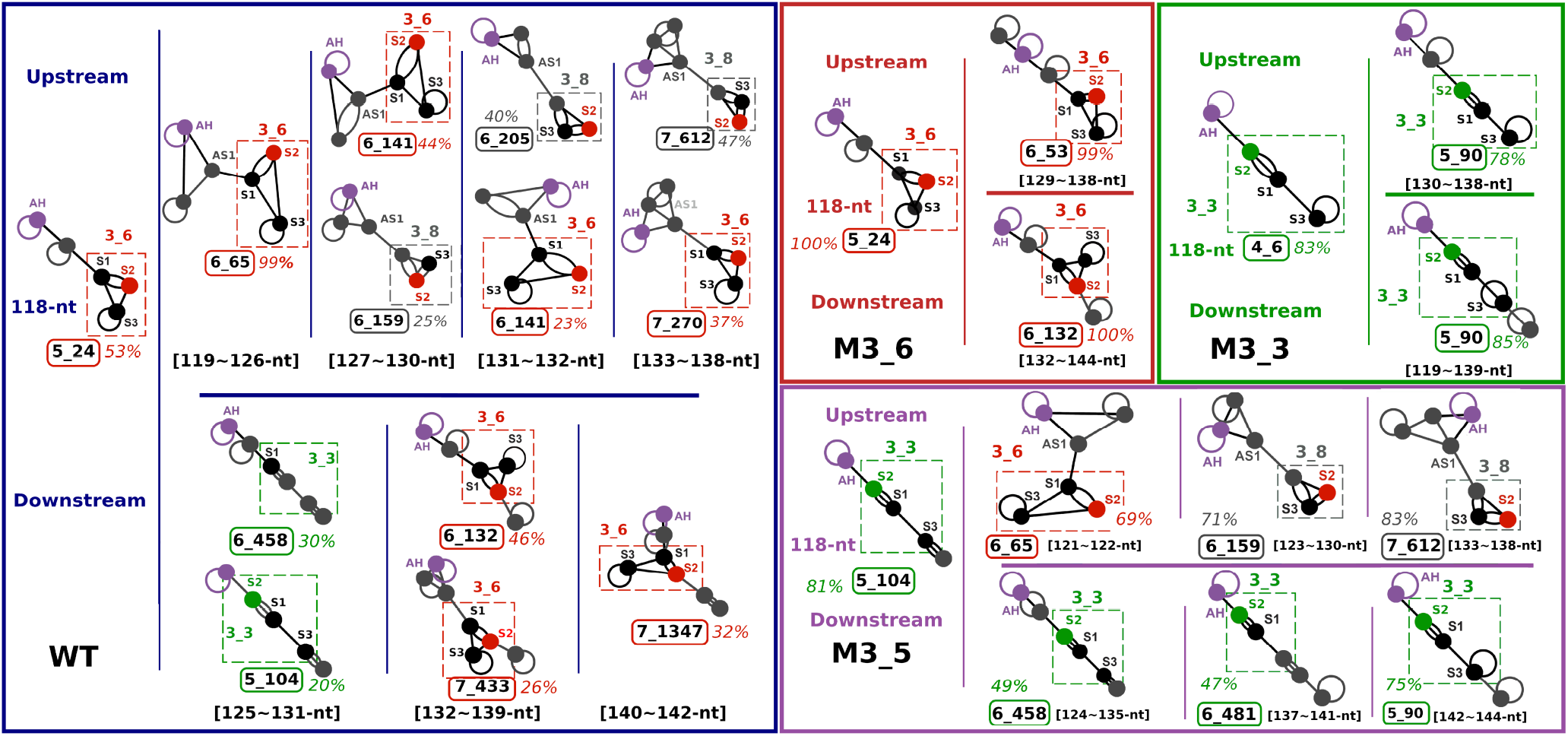
Dominant motifs for the FSE wildtype and mutant systems in this work with additional downstream and upstream sequence. Dual graphs of the dominant motif at certain sequence lengths are shown with AH labeled in purple and central FSE motif in boxes. For each system, the dominant motif is shown for the 118-nt base sequence on the left. Dominant motifs from the landscapes with additional downstream sequence are on the top, and those from upstream landscapes are on the bottom. The FSE subgraphs are labeled in red for 3_6, green for 3_3, gray for 3_8. Whole AH-FSE motifs are labeled with the highest percentage in the noted sequence range and boxed in colors according to the same grouping from the landscapes in Figure 2

As we add downstream residues to the RNA, we see that most of the resulting motifs for the FSE include a stable AH with a downstream 3_6 pseudoknot until the sequence reaches 125-nt (i.e., 7 downstream residues added); then, the majority of the motifs adopt a 3_3 FSE pseudoknot instead. The exact dual graphs obtained at different lengths are shown in Figure 3. For example, at 125-nt, the AH and 3_3 containing fold is a 5_104 dual graph motif. Figure 4 illustrates the shift from AH and 3_6-containing fold to AH and 3_3-containing fold as red to green stems, where S2 reforms at longer sequences.

**Fig. 4.**
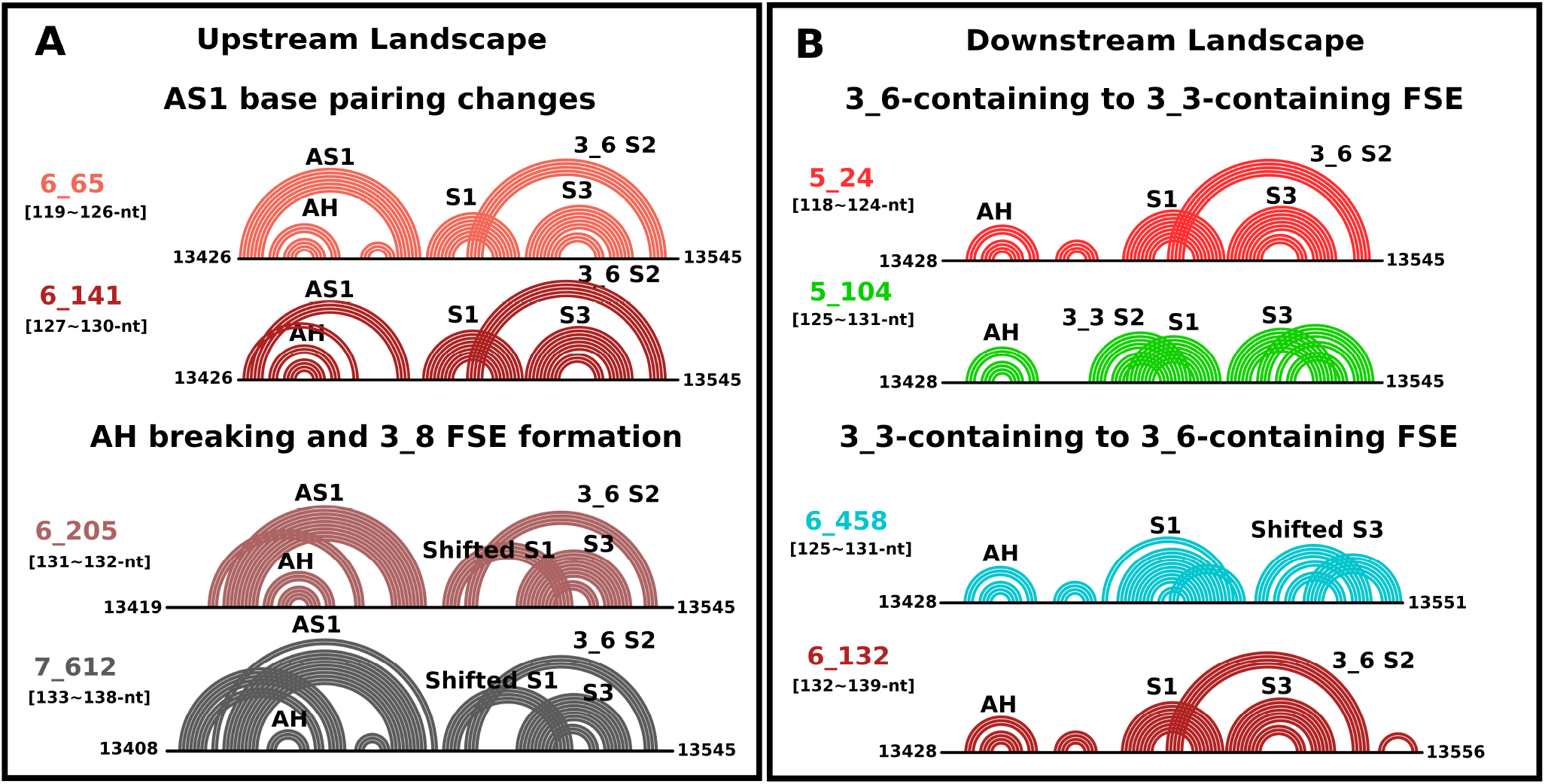
Transitions of 2D folding from the dominant motifs of FSE wildtype in this work with additional upstream (left) and downstream (right) sequence are shown in arc plots. 3_6 FSE containing motifs are colored in red; similarly, 3_3 FSE containing motifs are in green, and 3_8-containing motifs are in gray.

When sequences reach lengths above 132-nt, motifs 6_132 and 7_1347 (dual graphs in Figure 3) combine AH and a stable 3_6 FSE. Both motifs contain the hairpin downstream of the AH that blocks the formation of 3_3 S2, triggering a transition back to 3_6.

As we add upstream residues to the wildtype FSE, an Alternative Stem 1 (AS1) upstream forms, protecting the AH and blocking formation of 3_3 S2 and 3_5 S2 (Figure 4, right). This favors 3_6 topologies as well as a 3_8 pseudoknot (dual graphs in Figure 3 and related 2D folds in Figure 4, left). The 3_8 pseudoknot (as first reported in (52)) has three stems intertwined without a self-loop (motifs with 3_8 are colored in dark gray in Figures 2 and 4). It can form from a 3_6 pseudoknot with a shifted S1 (in this work) or S3 [59].

As clearly seen in Figure 4, right, AS1’s increasing size can block or alter S1. Without S1, “other” motifs and motifs including the AH and 3_8 can form, colored in light and dark gray (Figure 2).

To investigate the folding of the AH-FSE region more thoroughly, especially AS1 formation at long sequences, we also extend symmetrically in Figure 2B the 144-nt symmetric construct we previously developed, which includes 30-nt upstream and downstream of the central FSE region. We focus on a 156-nt construct, which contains the long AS1. In this scenario of longer upstream sequence, the potential 3_3 FSE changes to 3_6, and the 3_6 pseudoknot also faces competition from other folds. Many of these folds have 3_8 pseudoknot-containing motifs, which uses S2 and S3 of 3_6 and a short shifted S1.

### Comparison Between Our Landscape Predictions and Published DMS Chemical Probing Experiments

We compare our predicted structures with DMS-MaP results by Pekarek et al. (36) for the most comparable sequence/construct. Our 143-nt construct with 34 residues upstream of the slippery site and 25 residues downstream of the 77-nt central FSE aligns well with the 141-nt FSE-V2 construct (33) (only 2-nt difference at the 5^′^-end). In Figure 5A, we show consistent dominant 3_6 FSE predicted in our landscape in agreement with the findings of Pekarek et al. (36). The differences include the minor 3_3 conformation predicted in our landscape (21.67% at 143-nt) and the small hairpin following AH in the 3_6-containing motifs (see complete prediction profile in Figure S2). The dominant 3_6-containing motif for the AH-FSE region predicted in our landscape is conserved from shorter sequences, such as 118-nt, to sequences that match with experiments in (36).

**Fig. 5.**
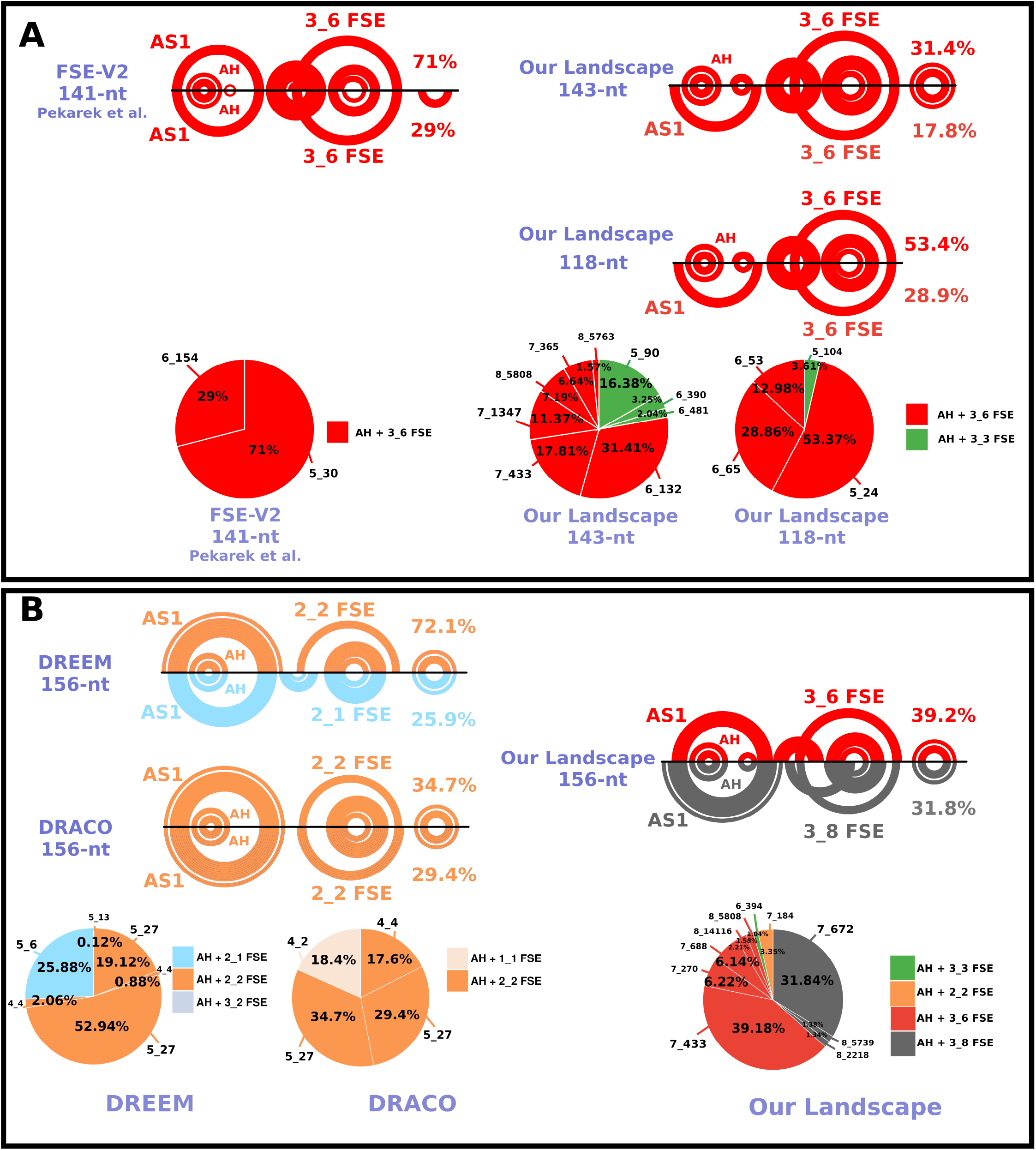
Comparison of our landscape predictions with DMS-MaP chemical probing supported structures. Two folds for each sequence and prediction combination with the largest fraction are shown in arc plots. (A) Comparison of 143-nt from wildtype downstream landscape and 141-nt FSE-V2 construct by Pekarek et al. (36). (B) Comparison of 156-nt from the wildtype long sequence landscape and the same construct from DMS-MaPseq chemical probing experiment followed by DREEM (46) and DRACO (48) clustering and ShapeKnots (47) predictions. The overall distributions of motifs for DREEM, DRACO and our landscapes are shown in a pie chart format, and full details of the graph motifs are given in Figure S2.

### 156-nt DMS Data and Computational Landscapes Assessment

To assess our computational predictions, we also perform DMS chemical probing experiments on 156-nt constructs (see Methods) and analyze the information using DREEM (46) and DRACO (48) (see Figure 5B and Figure S3). Sequencing reads are clustered and the reactivity profiles guide RNA predictions via ShapeKnots (47). For the 156-nt DMS data analyzed here, dominant folds contain a long AS1, AH and 2_2 FSE, with S1 absent but S2 and S3 of 3_6 present. S1 cannot form because of the extra base pairs (AU and GC pairs) in AS1 (in Figure 5B). DREEM also predicts an alternative 2_1 FSE, where FSE S1 and S3 persist without S2. In comparison, our computed landscapes for this length in Figure 2B predict a minority of 2_2 (gray, 3.35%), 49.11% AH-containing 3_6 motif (8-bp AS1), and 33.22% 3_8 containing motif with shifted S1 and 12-bp AS1.

All these motifs are consistent with the mechanistic picture emerging from our landscape (next section) and the fact that DREEM and DRACO programs do not favor formation of pseudoknots at this long sequence length.

In our landscape, as sequence length increases from 141 to 156-nt, the distribution of different motifs (mainly 3_6 FSE in combination with an 8-bp AS1 and 3_8 FSE in combination with a 12-bp AS1) changes consistently with the increasing fraction of motifs with a 12-bp AS1. In these motifs, S1 is disrupted, consistent with DMS data, confirming that S1 can form only when AS1 is shorter or equal to 10-bp, though the 3_8 pseudoknot with a shifted short stem may also emerge.

The reliability of our NUPACK landscapes can be determined by examining the predictions on 77-nt wildtype and motif strengthening mutants (in Table S2). For long sequences, such as 156-nt, 3_6 FSE is also predicted by other softwares (Figure S4) despite different base pairs variations for the AS1 region. The DMS-guided predictions via DREEM and DRACO also exhibit similar behavior. DREEM and DRACO agree with NUPACK at short sequences, such as 77-nt, but inconsistencies quickly arise for longer RNAs, such as 87-nt and 114-nt, as we show in Table S3. Our close examination make clear that caution is warranted when interpreting the results for both pure computational and chemical probing guided predictions at long sequences, especially regarding pseudoknots, which tend to be underpredicted by softwares such as DREEM, DRACO and DANCE-MaP (37).

Thus, together, our landscapes and DMS probing data all agree that this region of the SARS-CoV-2 genome adopts multiple, competing conformations that are part of the conformational landscape and whose relative proportions are highly sensitive to the RNA length and to analysis programs. Our landscape makes clear that 3_6 S2 can coexist with AH and that an AS1 longer than 10-bp results in shifted S1. Without S1, stem-loops (2_1 or 2_2) observed in multiple experimental studies (53; 30; 52) and in long constructs by our DMS-MaPseq (156-nt, Figure 5B, and 222-nt FSE-containing sequence, Figure S5) are favored instead of the pseudoknot 3_6.

### Length-dependent Folding Mechanism of AH-FSE Sequence from Conformational Landscapes

Combining our conformational landscape information and experimental findings, we propose in Figure 6 that the length-dependent FSE folding might follow this trajectory as the ribosome unwinds during translation. Namely, as the ribosome moves along the wildtype sequence (upstream sequence decreasing), our landscapes in Figures 2 reveal the following folding mechanisms of the AH-FSE template region:

**Fig. 6.**
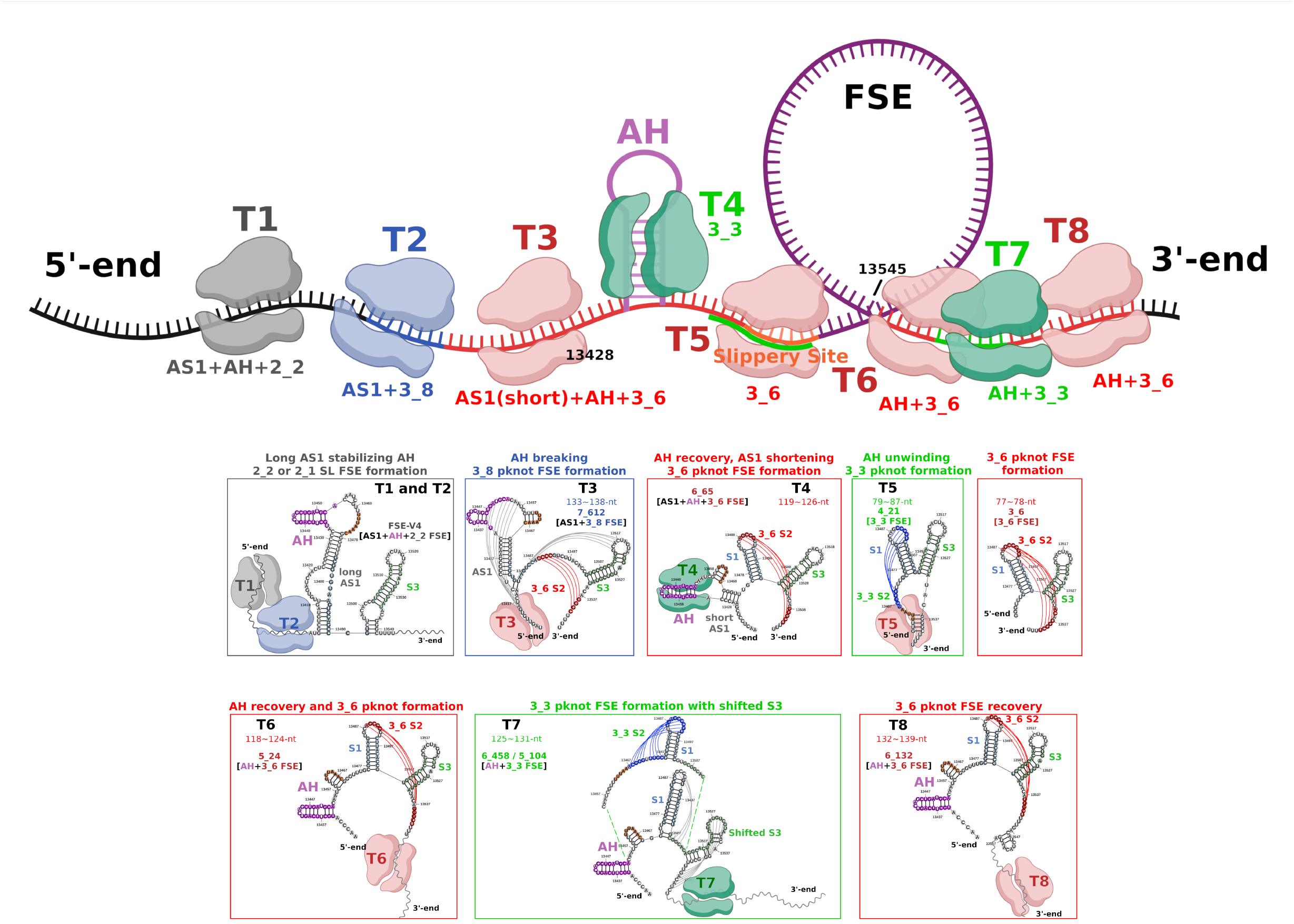
Proposed FSE conformational transition resulting from the unwinding by ribosomes during translation. In our top schematic diagram, the topology of central FSE region changes as the ribosome moves along the RNA genome. Ribosomes located at timesteps T1 to T8 are colored based on the associated FSE fold motif, red for 3_6 pseudoknot, green for 3_3 pseudoknot, and gray for unknotted stem-loop 2_2. The RNA templates are boxed and colored to match the ribosome positions. The arrows indicate the range of ribosome unwinding of the local RNA. Folds of the AH-FSE region are shown in secondary structure visualization via RNAcanvas (59).

1. Long sequences promote the formation of unknotted stem-loops that contain an elongated AS1 of 12 base pairs or more and either a 2_2 (simple stem-loop) or 2_1 FSE **(template 1)**.
2. Disruption of partial AS1 results in a shifted S1 in the predicted 3_8 motif (for central FSE) **(template 2)**.
3. Further unwinding leads to a shortened AS1, AH and 3_6 FSE **(template 3)**.
4. Unwinding of AH results in 3_3 FSE formation when the slippery site is available for pairing **(template 4)**.
5. When the ribosome’s movement renders the slippery site inaccessible, the 3_6 FSE is reinstated **(template 5)**.
6. Additional downstream residues from the 77-nt central FSE support a 3_3 FSE (see 5_109, for example) **(template 6 to 7)**.
7. As the ribosome further moves away from FSE, 3_6 FSE is reestablished, alongside AH **(template 8)**.

In general, short stems, for example, 3_3 S2 and 3_8 shifted S1, can form temporarily after adding sequence upstream or downstream, while long stems, such as 2_2 stems and S3, are more stable. In a separate work, we simulate the 3_3 to 3_6 pseudoknot transition, confirming its importance on aligning the FSE region in the narrow mRNA channel (54).

### M3_3 and M3_6 SARS-CoV-2 Conformational Landscape

In our previous work (25), we designed a series of pseudoknot-strengthening mutants to fold onto the desired FSE conformation at 77-nt. Here, we extend our studies and calculations to investigate whether these mutants retain the designed folds for longer sequence contexts, when AH is considered.

Our M3_6 folds almost perfectly into a stable AH and a downstream 3_6 FSE (Figure 7), both when adding downstream or upstream residues. Additional hairpins are appended downstream or upstream of the AH-FSE region (Figure S6A).

**Fig. 7.**
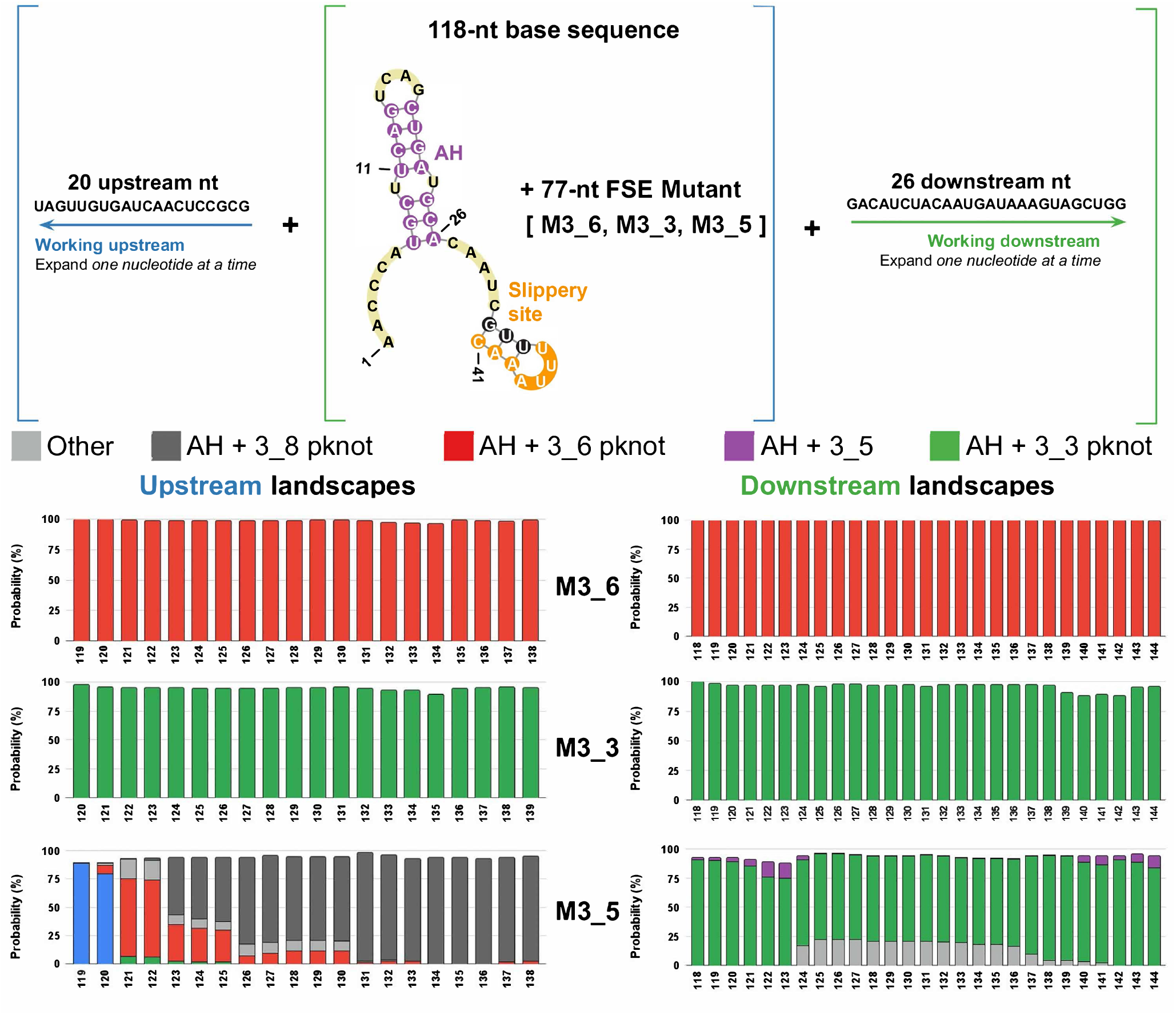
Conformational landscapes of our three motif strengthening mutants M3_6, M3_3, and M3_5 with expanding sequence upstream and downstream. Motifs containing 3_6 and AH are in red; motifs containing 3_3 and AH are in green; and motifs containing 3_5 and AH are in purple. See Figure 3 for examples of folds at different lengths.

In contrast to the wildtype variant, M3_3 fares well in folding into both an upstream AH and a downstream 3_3 FSE both when adding downstream or upstream residues (for example, 5_90 in Figure 3). Adding upstream residues generates a stable AH along with a stable downstream 3_3 FSE. However, S3 can be replaced by downstream hairpins, e.g., 6_390, after appending downstream residues to M3_3 at 140∼142-nt (shown in Figure S6B).

### M3_5 SARS-CoV-2 Conformational Landscape

The conformational landscapes reveal that the M3_5 mutant refolds as downstream residues are added (Figure 7). Many of the resulting motifs hold a 3_3 FSE instead of the expected 3_5 FSE, underscoring the minor role of the three-way junction fold in the conformational ensemble. A possible AH and 3_5-containing motif is 6_52 at 140∼144-nt for 5-10% shown in Figure S7. The formation of 3_3 S2 in 6_458 (Figure S6C) is favored over that of 3_5 S2 in 6_52 for most of the motifs.

Similar to the wildtype variant, many of these motifs possess AS1 after adding upstream residues, preventing 3_5 S2 from forming. A motif with AS1 intertwined with a shifted 3_5 S2 exists at 122∼126-nt very rarely for 2-6% (Figure S1).

### M3_5 SARS-CoV-2 RAG-IF Mutation Design for 3_5 at Long Sequences

To construct a more stable 3_5 FSE for long sequences, we utilize our mutation design algorithm, RAG-IF (see **Methods**), to search for minimal mutations.

We focus on the downstream residues of 3_3 S2, hoping to simultaneously break 3_3 S2 and block 3_6 S2 to make 3_5 S2 favorable. We ran RAG-IF for each motif that did not have a 3_5 FSE, hoping to find mutations that would cause a motif to maintain AH and a stable downstream 3_5 FSE. After mutating all eight motifs, we looked for commonalities among the many possible mutation sequences for each motif.

Ultimately, we found seven total mutations: [G16U, U17G, G21A, C23U, C24A, C47A, G52U] (Figure 8A, bottom). Specifically, C23U and C24A break two G-C pairs in 3_3 S2, weakening 3_3 S2 and making the formation of 3_5 S2 much more favorable. The three mutations G21A, C23U, and C24A strengthen 3_5 S1 and block the formation of 3_6 S2. Together, these mutations break 3_3 S2, block the formation of 3_6 S2, and strengthen 3_5 FSE.

**Fig. 8.**
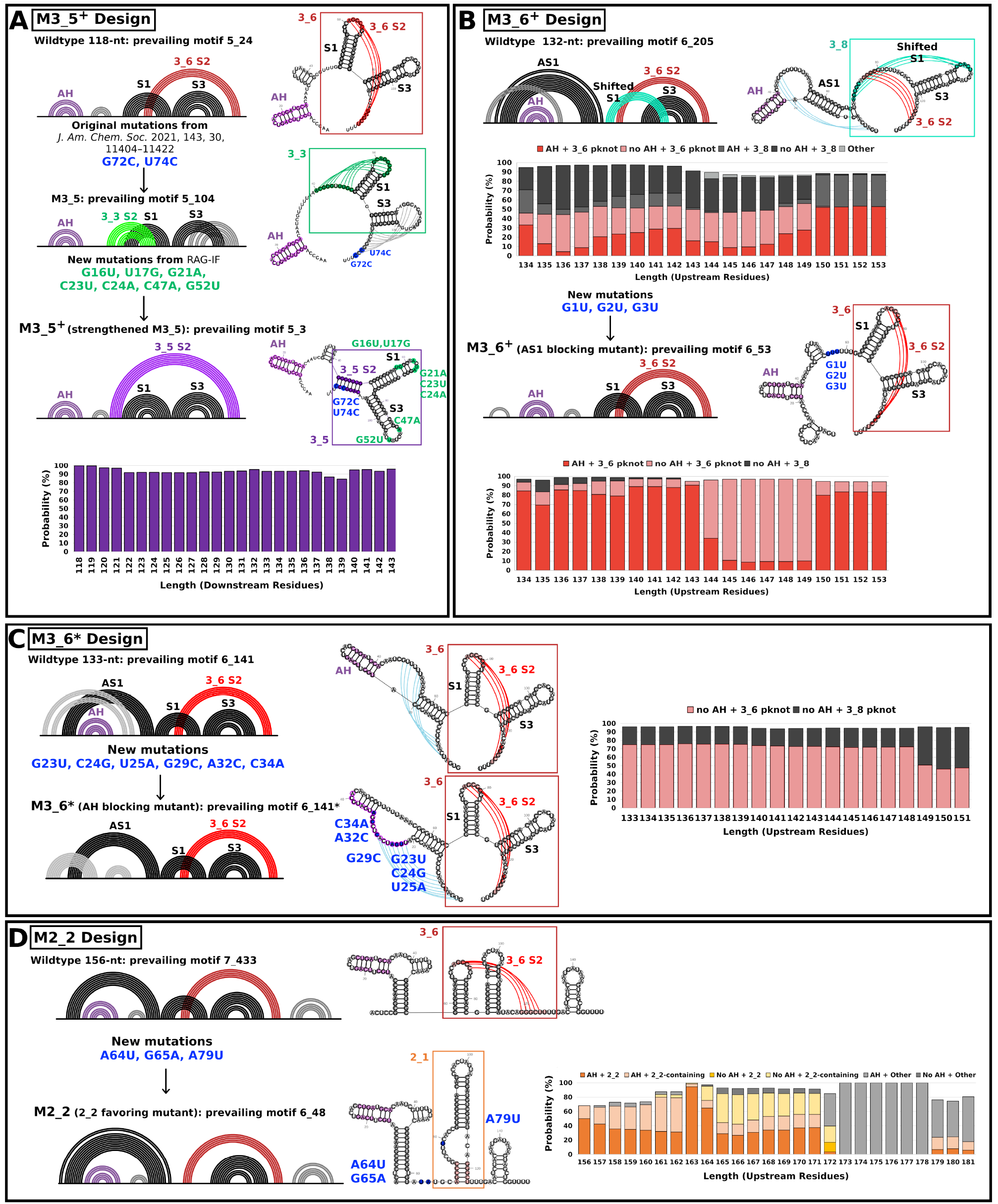
Mutant designs to stabilize 3_5 (M3_5^+^), block AS1 (M3_6^+^), block AH (mutant M3_6*), or favor stem-loop 2_2 by blocking S1 (M2_2) in long FSE-containing sequences. (A) To design the mutant (M3_5^+^) that supports 3_5 junction formation, we use RAG-IF and multiple 2D structure prediction program screening to determine the minimal mutations. The original mutations in M3_5 (25) are denoted in blue. To stabilize 3_5 for long sequence contexts, we identify 7 mutations by RAG-IF and add to M3_5. New mutations from this work are denoted in green. The conformational landscapes are calculated for each mutant to examine motif stability under a long sequence context. (B) To block AS1 and form the 3_6 pseudoknot, we manually mutate G1 to G3 in the 77-nt FSE region to U (denoted in blue) and examine the folding for 132-nt and stability in long RNAs (expanding upstream from 134 to 153-nt). Three G-C pairs in the 12 bp AS1 at 132-nt are disrupted, and thus AS1 with 3_8 is blocked in our resulting M3_6^+^ mutant. (C) To block AH and form the 3_6 pseudoknot, we use RAG-IF and 2D structure prediction programs IPknot and NUPACK to screen and determine the minimal mutations. For the 133-nt FSE-containing sequence, we mutate the AH region and replace AH with a stem knotted with AS1 and a small hairpin. The conformational landscape expanding upstream shows the stability of the mutant motif. (D) To block S1 and favor the 2_2 stem-loop, we design the sequence with minimal mutations for the 156-nt FSE-containing sequence. Three mutations are introduced to destroy three base pairs (2 AU pairs and 1 GC pair) in S1 and thus favor the 2_2 stem-loop. The conformational landscape expanding upstream confirms our target stem-loop stability.

When examining the motifs created from this new mutation sequence (M3_5^+^), the seven mutations clearly help stabilize 3_5 FSE over a wide range of lengths (Figure 8A).

### AS1 or AH Blocking Mutation Design for 3_6 at Long Sequences

To stabilize the pseudoknot formation for long RNA sequences, and further validate our mechanistic landscape (Figure 6), we mutate the FSE region to block the formation of AS1. From Figure 5 we know that AS1 is strengthened when S1 residues base pair with upstream sequence. A 12-bp AS1 further prevents the formation of S1 and 3_6 pseudoknots. Thus, we mutate residues G1, G2 and G3 in S1 region to U, blocking AS1 G-C pairs. In Figure 8B, we examine the folding of the mutated sequence for 132-nt, where a 12-bp AS1 exists for the wildtype. The G-U mutations indeed block AS1 formation and recover S1 in the 3_6 pseudoknot. We expand the sequence with additional upstream residues and see that the 3_6 FSE replaces folds with both AS1 and 3_8 and thereby dominates the landscape with high stability as designed. We call this mutant M3_6^+^.

Alternatively, we stabilize the 3_6 pseudoknot by destroying AH through mutations in the hairpin region while maintaining AS1 and the 3_6 pseudoknot (M3_6*). From Figure 8C, we split AH strands into two stems, including an 8-bp stem knotted with AS1 and a 5-bp hairpin. Thus, the upstream stem-loop seen in our 156 and 222-nt FSE-containing sequences in Figure 5 and Figure S5 and the literature (30; 36) switches to a 3_6 pseudoknot. As AS1 expands, the 3_8 motif as seen in the wildtype system also emerges as part of the landscape.

### M2_2 Stem-loop Favoring Mutation Design for Stem-loop at Long Sequences

Guided by our mechanism in Figure 6, we block the formation of S1 to favor the stem-loop by mutatating the FSE S1 region. Three mutations can disrupt S1 and support the 2_2 stem-loop for our M2_2 mutant in (Figure 8D). As S1 is inhibited, AS1 extends from 8-bp to 12-bp, which is consistent with the observations from our landscape. The 2_2 and 2_2-containing motifs are stable up to 171-nt. Upstream sequences further extend AS1 beyond 171-nt, which disrupts the stem-loop in the central FSE region. This automatic mutation design via RAG-IF confirms that disrupting S1 results in a 12-bp AS1 and 2_2 FSE, supporting our findings of the effect of upstream sequence on the FSE pseudoknot and stem-loop conversion.

Combining the findings from this study with those of earlier research (24; 25), the robust picture as shown in Figure 6 emerges of how AH and AS1 compete/interfere with the short sequence FSE pseudoknot 3_6.

## Discussion

To elucidate the length and context dependent RNA folding and its impact on RNA function, we have developed conformational landscapes for different folding/refolding scenarios, considering that RNA can adopt multiple conformations at a given length. Computational predictions used in these landscapes offer a fast way to explore the folding space, when experimental characterization for a large number of RNA sequences of different lengths is not available/feasible. However, such imperfect predictions depend on the methodology (energy-based, covariation-based, thermodynamic and comparative analysis integrated, traditional machine learning-based and deep learning-based, see review (55)), and are not easily comparable to one another. Moreover, not all programs compute suboptimal folds or conformational weights, as required for our landscapes. The accuracy of predictions is sequence length-dependent. For example, some programs correctly predict the 77-nt FSE and motif stabilizing mutants (Table S2) but fail to predict the 7-bp AH for long sequences (Figure S4). Chemical probing data constraints can guide these RNA 2D predictions, but different approaches are needed to process the experimental data and analyze multiple conformational states. Newer pipelines (46; 48) that incorporate experimentally determined structural data into computational modeling can help analyze RNA structures, but because of disparate clustering capabilities and prediction sensitivity on the experimental reactivity profile, these methods may also produce divergent outcomes (as in Figure 5). One remaining challenge in reconciling these different analyses is the lack of a reference experimental approach for determining alternative conformational distributions in large (over 150-nt) RNAs.

Taking into account the limitations of various computational pipelines in producing alternative clusters, predicting pseudoknots, and integrating SHAPE/DMS chemical probing data, it is not straightforward to adhere to a single program or methodology. Functional RNA structures, such as the frameshift element, are known to adopt folds that depend sensitively on specific cellular and biological context. While much caution is warranted in creating and interpreting the various conformational landscapes, it is useful to try to combine all the clues we have reported here into a complex dynamic view of the frameshifting RNA region. In general, resolving multiple conformational landscapes for RNA is difficult, and different computational platforms do not always lead to consensus. However, our various conformational landscapes as presented here together with experimental data yield a robust picture overall.

For SARS-CoV-2 FSE, the 3_6 pseudoknot is prevalent at short to medium RNA lengths (26; 27; 2; 29), with 3_3 appearing too at medium lengths, and the two forms interchange in short to medium lengths (25). When the sequence immediately downstream of the central 77-nt FSE is included in the folding, 3_3 emerges. The 3-way junction 3_5 also appears as a very minor player in specific contexts (25). Over extended lengths, alternative folds increasingly replace the 3_6 and 3_3 pseudoknots as summarized in our proposed mechanism in Figure 6. Namely, with the emergence of a long, stable AS1, S1 is shortened and displaced, and FSE takes on a 3_8 pseudoknot fold, where S1 is intertwined with both S2 and S3. Other simpler folds may also be favored such as stem-loops 2_2 (no S1) and 2_1 (with S1). AH is compatible at low sequence lengths with pseudoknots 3_6 and 3_3 when AS1 does not form. As AS1 develops at medium lengths, it replaces S1 and leads to a shifted S1 in 3_6 or 3_8 along with AH. With more additional upstream and downstream sequence, the anchoring helix AS1 expands and blocks FSE pseudoknot formation. For some long lengths, AH can unfold so that the AH residues contribute to a knotted stem upstream of AS1. When the ribosome unwinds the long anchoring helix AS1, the downstream sequence can refold back into the 3_6 pseudoknot.

The AH can also form without AS1, perhaps explaining why the hairpin remains even though the sequence is less conserved in the upstream region. Its formation is also independent of downstream FSE region, whereas AS1 involving the spacer sequence and S1 sequence has a large impact on downstream FSE. This is consistent with the observation that the spacer sequence plays a significant role in frameshifting efficiency (2; 56). A longer AS1 likely re-forms as a consequence of refolding when the 3_3 or 3_6 pseudoknot unwinds and the translating ribosome is away from the ORF1a,b overlap region.

From the literature (17), AH has been shown to trigger ribosome dissociation from the mRNA and thus decrease the fraction of ribosomes available to frameshifting. We further propose in Figure 6 that when the ribosome remains interacting with the RNA, it unwinds AH and possibly causes the downstream FSE to explore various folds on the altered RNA free energy landscape, mainly 3_6 and 3_3 pseudoknots. Thus, structural plasticity might play a key role in frameshifting, as we highlight in a separate work that follows the 3_3 to 3_6 pseudoknot transition (54). Moreover, the pseudoknot stability/compactness is related to the frameshifting efficiency (17; 57). When the ribosome passes the overlap region, AH re-forms and pseudoknots 3_6 or 3_3 can be restored, until stem-loops replace the pseudoknot when the ribosome is distant from the overlap region.

For the motif strengthening mutant systems, our mutants M3_3 and M3_6 are quite stable at varying lengths, and coexist with AH while formation of long AS1 is blocked. This indicates that our mutation designs in Ref (25) effectively strengthen the pseudoknots and block conformational transitions that result from translating ribosomes. Experimental frameshifting efficiencies support these findings (37; 36).

The M3_5 exhibits a dominant 3_3 when adding upstream residues, and because the 3_3 S2 is stronger than 3_5 S2, 3_5 cannot dominate. With more upstream residues included, AS1 blocks the formation of 3_5 S2 and S1 in M3_5. To restore 3_5, our designed mutant M3_5^+^ here blocks 3_3 and maintains 3_5 for longer sequence. Our new mutations in the 3_3 3^′^-strand break 3_3 S2 pairing and strengthen S1. Our M3_6^+^ mutant designed here blocks AS1 altogether and favors 3_6. The M3_6* mutant designed here disrupts AH base pairs and results in a 3_6 pseudoknot containing AS1. The M2_2 mutant blocks S1 and supports a stem-loop FSE in combination with a 12-bp AS1. These four mutants (M3_5^+^, M3_6^+^, M3_6*, and M2_2) that favor 3_5, 3_6, or 2_2 while destroying AS1, AH or S1 reinforce our mechanistic understanding in Figure 6 of the evolving template-driven folding/unfolding events and may be useful for antiviral approaches targeting a specific conformer. Their effect on frameshifting can be further examined by experiments.

Because each virus has evolved to create optimal rates of −1 PRF by balancing more and less stable structural features (17), alternative structures play an important role in frameshifting and viral propagation. Their formation depends on translation rates between conformers and ribosomal pausing (58). Forming a stable and unwinding-resistant conformation to impede translating ribosomes may be valuable for developing efficient frameshifting elements (58), as in AH protected by a stable AS1.

Overall, our computational work and experimental validations (25; 36) suggest an interactive model of AH-FSE structure modulated by translating ribosomes during frameshifting and largely influenced by AS1 formation. Blocking AS1 in our M3_6^+^ mutant here may allow more complex folds to form (Figure 8B). Additional experimental and computational perspectives will undoubtedly shed further insight into this complex process.

## Author contributions statement

T.S. and A.L. conceived the computations and experiments, S.L., S.Y. and A.D. conducted the computations and experiments, S.L., S.Y., A.D., A.L. and T.S. analysed the results. S.L., S.Y., A.D., A.L. and T.S. wrote and reviewed the manuscript.

## Acknowledgments

We thank Shereef Elmetwaly for technical assistance and the support from the Simons Foundation through the NYU Simons Center for Computational Physical Chemistry.

We gratefully acknowledge funding from the National Science Foundation RAPID Award 2030377 from the Division of Mathematical Science and the Division of Chemistry, National Science Foundation Award DMS-2151777 and CHE-2330628 from the Division of Mathematical Sciences, National Institutes of Health R35GM122562 Award from the National Institute of General Medical Sciences, and Philip-Morris International to T. Schlick; R35GM140844 Award from National Institutes of Health to A. Laederach; Dean’s Undergraduate Research Fund from NYU College of Arts and Science to S. Lee; and Ramalingaswami Re-entry Fellowship (BT/RLF/Re-entry/02/2021) from Department of Biotechnology, Govt. of India to A. Dey.

## References

1. Y. Sun, L. Abriola, R. O. Niederer, S. F. Pedersen, M. M. Alfajaro, V. S. Monteiro, C. B. Wilen, Y.-C. Ho, W. V. Gilbert, Y. V. Surovtseva, et al., “Restriction of SARS-CoV-2 replication by targeting programmed - 1 ribosomal frameshifting,” Proc. Natl. Acad. Sci. U.S.A., vol. 118, no. 26, p. e2023051118, 2021.

2. P. R. Bhatt, A. Scaiola, G. Loughran, M. Leibundgut, A. Kratzel, R. Meurs, R. Dreos, K. M. O’Connor, A. McMillan, J. W. Bode, et al., “Structural basis of ribosomal frameshifting during translation of the SARS-CoV-2 RNA genome,” Science, vol. 372, pp. 1306–1313, 2021.

3. C. Varricchio, G. Mathez, T. Pillonel, C. Bertelli, L. Kaiser, C. Tapparel, A. Brancale, and V. Cagno, “Geneticin shows selective antiviral activity against SARS-CoV-2 by interfering with programmed -1 ribosomal frameshifting,” Antiviral Res., vol. 208, p. 105452, 2022.

4. Munshi, K. Neupane, S. M. Ileperuma, M. T. J. Halma, J. A. Kelly, C. F. Halpern, J. D. Dinman, S. Loerch, and M. T. Woodside, “Identifying inhibitors of -1 programmed ribosomal frameshifting in a broad spectrum of coronaviruses,” Viruses, vol. 14, Feb. 2022.

5. M. Yang, F. P. Olatunji, C. Rhodes, S. Balaratnam, K. Dunne-Dombrink, S. Seshadri, X. Liang, C. P. Jones, S. F. J. Le Grice, A.R. Ferré-D’Amaré, and J. S. J. Schneekloth, “Discovery of Small Molecules Targeting the Frameshifting Element RNA in SARS-CoV-2 Viral Genome,” ACS Med. Chem. Lett., vol. 14, pp. 757–765, 2023.

6. S.-H. Huang, S.-C. Chen, T.-Y. Wu, C.-Y. Chen, and C.-H. Yu, “Programmable modulation of ribosomal frameshifting by mRNA targeting CRISPR-Cas12a system,” iScience, vol. 26, p. 108492, 2023.

7. J. A. Kelly, M. T. Woodside, and J. D. Dinman, “Programmed -1 Ribosomal Frameshifting in coronaviruses: A therapeutic target,” Virology, vol. 554, pp. 75–82, 2021.

8. I. Brierley, P. Digard, and S. C. Inglis, “Characterization of an efficient coronavirus ribosomal frameshifting signal: requirement for an RNA pseudoknot,” Cell, vol. 57, no. 4, 1989.

9. S. Napthine, J. Liphardt, A. Bloys, S. Routledge, and I. Brierley, “The role of RNA pseudoknot stem 1 length in the promotion of efficient -1 ribosomal frameshifting,” J. Mol. Biol., vol. 288, pp. 305–320, 1999.

10. H. Kontos, S. Napthine, and I. Brierley, “Ribosomal Pausing at a Frameshifter RNA Pseudoknot Is Sensitive to Reading Phase but Shows Little Correlation with Frameshift Efficiency,” Mol. Cell. Biol., vol. 21, pp. 8657–8670, 2001.

11. P. V. Baranov, C. M. Henderson, C. B. Anderson, R. F. Gesteland, J. F. Atkins, and M. T. Howard, “Programmed ribosomal frameshifting in decoding the SARS-CoV genome,” Virology, vol. 332, pp. 498–510, 2005.

12. E. P. Plant, G.C. Pérez-Alvarado, J. L. Jacobs, B. Mukhopadhyay, M. Hennig, and J. D. Dinman, “A Three-Stemmed mRNA Pseudoknot in the SARS Coronavirus Frameshift Signal,” PLoS Biol., vol. 3, p. e172, 2005.

13. O. Namy, S. J. Moran, D. I. Stuart, R. J. C. Gilbert, and I. Brierley, “A mechanical explanation of RNA pseudoknot function in programmed ribosomal frameshifting,” Nature, vol. 441, pp. 244–247, 2006.

14. C.-P. Cho, S.-C. Lin, M.-Y. Chou, H.-T. Hsu, and K.-Y. Chang, “Regulation of Programmed Ribosomal Frameshifting by Co-Translational Refolding RNA Hairpins,” PLoS One, vol. 8, p. e62283, 2013.

15. H.-T. Hu, C.-P. Cho, Y.-H. Lin, and K.-Y. Chang, “A general strategy to inhibiting viral -1 frameshifting based on upstream attenuation duplex formation,” Nucleic Acids Res., vol. 44, no. 1, pp. 256–266, 2015.

16. M.-C. Su, C.-T. Chang, C.-H. Chu, C.-H. Tsai, and K.-Y. Chang, “An atypical RNA pseudoknot stimulator and an upstream attenuation signal for -1 ribosomal frameshifting of SARS coronavirus,” Nucleic Acids Res., vol. 33, pp. 4265–4275, 2005.

17. E. P. Plant, R. Rakauskaite, D. R. Taylor, and J. D. Dinman, “Achieving a Golden Mean: Mechanisms by Which Coronaviruses Ensure Synthesis of the Correct Stoichiometric Ratios of Viral Proteins,” J. Virol., vol. 84, pp. 4330–4340, 2010.

18. H. H. Gan, S. Pasquali, and T. Schlick, “Exploring the repertoire of RNA secondary motifs using graph theory; implications for RNA design,” Nucleic Acids Res., vol. 31, pp. 2926–2943, 2003.

19. M. Zahran, C. Sevim Bayrak, S. Elmetwaly, and T. Schlick, “RAG-3D: a search tool for RNA 3D substructures,” Nucleic Acids Res., vol. 43, pp. 9474–9488, 2015.

20. N. Baba, S. Elmetwaly, N. Kim, and T. Schlick, “Predicting Large RNA-Like Topologies by a Knowledge-Based Clustering Approach,” J. Mol. Biol., vol. 428, pp. 811–821, 2016.

21. T. Schlick, “Adventures with RNA graphs,” Methods, vol. 143, pp. 16–33, 2018.

22. G. Meng, M. Tariq, S. Jain, S. Elmetwaly, and T. Schlick, “RAG-Web: RNA structure prediction/design using RNA-As-Graphs,” Bioinformatics, vol. 36, pp. 647–648, 2020.

23. S. Jain, Y. Tao, and T. Schlick, “Inverse folding with RNA-As-Graphs produces a large pool of candidate sequences with target topologies,” J. Struct. Biol., vol. 209, no. 3, p. 107438, 2020.

24. T. Schlick, Q. Zhu, S. Jain, and S. Yan, “Structure-altering mutations of the SARS-CoV-2 frameshifting RNA element,” Biophys. J., vol. 120, pp. 1040–1053, 2021.

25. T. Schlick, Q. Zhu, A. Dey, S. Jain, S. Yan, and A. Laederach, “To Knot or Not to Knot: Multiple Conformations of the SARS-CoV-2 Frameshifting RNA Element,” J. Amer. Chem. Soc., vol. 143, pp. 11404–11422, 2021.

26. C. P. Jones and A.R. Ferré-D’Amaré, “Crystal structure of the severe acute respiratory syndrome coronavirus 2 (SARS-CoV-2) frameshifting pseudoknot,” RNA (Cambridge), vol. 28, no. 2, pp. 239–249, 2022.

27. C. Roman, A. Lewicka, D. Koirala, N. Li, and J. Piccirilli, “The SARS-CoV-2 programmed −1 ribosomal frameshifting element crystal structure solved to 2.09 Å using chaperone-assisted RNA crystallography,” ACS Chem. Biol., vol. 16, pp. 1469–1481, 2021.

28. A. Wacker, J. E. Weigand, S. R. Akabayov, N. Altincekic, J. K. Bains, E. Banijamali, O. Binas, J. Castillo-Martinez, E. Cetiner, B. Ceylan, et al., “Secondary structure determination of conserved SARS-CoV-2 RNA elements by NMR spectroscopy,” Nucleic Acids Res., vol. 48, pp. 12415–12435, Dec. 2020.

29. K. Zhang, I. N. Zheludev, R. J. Hagey, R. Haslecker, Y. J. Hou, R. Kretsch, G. D. Pintilie, R. Rangan, W. Kladwang, S. Li, et al., “Cryo-EM and antisense targeting of the 28-kDa frameshift stimulation element from the SARS-CoV-2 RNA genome,” Nat. Struct. Mol. Biol., vol. 28, pp. 747–754, 2021.

30. T. C. T. Lan, M. F. Allan, L. E. Malsick, J. Z. Woo, C. Zhu, F. Zhang, S. Khandwala, S. S. Y. Nyeo, Y. Sun, J. U. Guo, et al., “Secondary structural ensembles of the SARS-CoV-2 RNA genome in infected cells,” Nat. Commun., vol. 13, p. 1128, 2022.

31. I. Manfredonia, C. Nithin, A. Ponce-Salvatierra, P. Ghosh, T. K. Wirecki, T. Marinus, N. S. Ogando, E. J. Snijder, M. J. van Hemert, J. M Bujnicki, and D. Incarnato, “Genome-wide mapping of SARS-CoV-2 RNA structures identifies therapeutically-relevant elements,” Nucleic Acids Res., vol. 48, no. 22, pp. 12436–12452, 2020.

32. C. Iserman, C. A. Roden, M. A. Boerneke, R. S. G. Sealfon, G. A. McLaughlin, I. Jungreis, E. J. Fritch, Y. J. Hou, J. Ekena, C. A. Weidmann, et al., “Genomic RNA Elements Drive Phase Separation of the SARS-CoV-2 Nucleocapsid,” Molecular Cell, vol. 80, pp. 1078–1091.e6, 2020.

33. W. Sanders, E. J. Fritch, E. A. Madden, R. L. Graham, H. A. Vincent, M. T. Heise, R. S. Baric, and N. J. Moorman, “Comparative analysis of coronavirus genomic RNA structure reveals conservation in SARS-like coronaviruses,” bioRxiv, p. 2020.06.15.153197, 2020.

34. L. Sun, P. Li, X. Ju, J. Rao, W. Huang, L. Ren, S. Zhang, T. Xiong, K. Xu, X. Zhou, et al., “In vivo structural characterization of the SARS-CoV-2 RNA genome identifies host proteins vulnerable to repurposed drugs,” Cell, vol. 184, pp. 1865–1883.e20, 2021.

35. S. Yan, Q. Zhu, S. Jain, and T. Schlick, “Length-dependent motions of SARS-CoV-2 frameshifting RNA pseudoknot and alternative conformations suggest avenues for frameshifting suppression,” Nat. Commun., vol. 13, p. 4284, 2022.

36. L. Pekarek, M. M. Zimmer, A.-S. Gribling-Burrer, S. Buck, R. Smyth, and N. Caliskan, “Cis-mediated interactions of the SARS-CoV-2 frameshift RNA alter its conformations and affect function,” Nucleic Acids Res., vol. 51, no. 2, pp. 728–743, 2023.

37. A. Dey, S. Yan, T. Schlick, and A. Laederach, “Abolished frameshifting for predicted structure-stabilizing SARS-CoV-2 mutants: Implications to alternative conformations and their statistical structural analyses.” bioRxiv, Mar. 2024.

38. S. Pasquali, H. H. Gan, and T. Schlick, “Modular RNA architecture revealed by computational analysis of existing pseudoknots and ribosomal RNAs,” Nucleic Acids Res., vol. 33, pp. 1384–1398, 2005.

39. J. Gevertz, H. H. Gan, and T. Schlick, “In vitro RNA random pools are not structurally diverse: A computational analysis,” RNA, vol. 11, pp. 853– 863, 2005.

40. K. Sato, Y. Kato, M. Hamada, T. Akutsu, and K. Asai, “IPknot: fast and accurate prediction of RNA secondary structures with pseudoknots using integer programming,” Bioinformatics, vol. 27, pp. i85–i93, 2011.

41. J. N. Zadeh, C. D. Steenberg, J. S. Bois, B. R. Wolfe, M. B. Pierce, A. R. Khan, R. M. Dirks, and N. A. Pierce, “NUPACK: Analysis and design of nucleic acid systems,” Journal of Computational Chemistry, vol. 32, pp. 170–173, 2011.

42. R. Dirks and N. Pierce, “A partition function algorithm for nucleic acid secondary structure including pseudoknots,” J. Comput. Chem, vol. 24, pp. 1664–1677, 2003.

43. K. A. Wilkinson, E. J. Merino, and K. M. Weeks, “Selective 2′-hydroxyl acylation analyzed by primer extension (SHAPE): quantitative RNA structure analysis at single nucleotide resolution,” Nat. Protoc., vol. 1, pp. 1610–1616, Aug. 2006.

44. M. J. Smola, G. M. Rice, S. Busan, N. A. Siegfried, and K. M. Weeks, “Selective 2′-hydroxyl acylation analyzed by primer extension and mutational profiling (SHAPE-MaP) for direct, versatile and accurate RNA structure analysis,” Nat. Protoc., vol. 10, pp. 1643–1669, Nov. 2015.

45. S. Busan and K. M. Weeks, “Accurate detection of chemical modifications in RNA by mutational profiling (MaP) with ShapeMapper 2,” RNA, vol. 24, pp. 143–148, Feb. 2018.

46. P. J. Tomezsko, V. D. A. Corbin, P. Gupta, H. Swaminathan, M. Glasgow, S. Persad, M. D. Edwards, L. Mcintosh, A. T. Papenfuss, A. Emery, et al., “Determination of RNA structural diversity and its role in HIV-1 RNA splicing,” Nature, vol. 582, no. 7812, pp. 438–442, 2020.

47. C. E. Hajdin, S. Bellaousov, W. Huggins, C. W. Leonard, D. H. Mathews, and K. M. Weeks, “Accurate SHAPE-directed RNA secondary structure modeling, including pseudoknots,” Proc. Natl. Acad. Sci. U.S.A., vol. 110, pp. 5498–5503, 2013.

48. E. Morandi, I. Manfredonia, L. M. Simon, F. Anselmi, M. J. van Hemert, S. Oliviero, and D. Incarnato, “Genome-scale deconvolution of RNA structure ensembles,” Nat. Methods., vol. 18, no. 3, pp. 249–252, 2021.

49. J. Zhang, K. Kobert, T. Flouri, and A. Stamatakis, “PEAR: a fast and accurate Illumina Paired-End reAd mergeR,” Bioinformatics, vol. 30, pp. 614–620, Mar. 2014.

50. D. Incarnato, E. Morandi, L. M. Simon, and S. Oliviero, “RNA Framework: an all-in-one toolkit for the analysis of RNA structures and post-transcriptional modifications,” Nucleic Acids Res., vol. 46, p. e97, 2018.

51. M. Zubradt, P. Gupta, S. Persad, A. M. Lambowitz, J. S. Weissman, and S. Rouskin, “DMS-MaPseq for genome-wide or targeted RNA structure probing in vivo,” Nat. Methods., vol. 14, pp. 75–82, 2017.

52. N. C. Huston, H. Wan, M. S. Strine, R. de Cesaris Araujo Tavares, C. B. Wilen, and A. M. Pyle, “Comprehensive in vivo secondary structure of the SARS-CoV-2 genome reveals novel regulatory motifs and mechanisms,” Mol. Cell, vol. 81, pp. 584–598.e5, 2021.

53. C. Cao, Z. Cai, X. Xiao, J. Rao, J. Chen, N. Hu, M. Yang, X. Xing, Y. Wang, M. Li, et al., “The architecture of the SARS-CoV-2 RNA genome inside virion,” Nat. Commun., vol. 12, no. 1, p. 3917, 2021.

54. S. Yan and T. Schlick, “Heterogeneous and multiple conformational transition pathways between pseudoknots of the SARS-CoV-2 frameshift element,” 2024. In Preparation.

55. J. Zhang, Y. Fei, L. Sun, and Q. C. Zhang, “Advances and opportunities in RNA structure experimental determination and computational modeling,” Nat. Methods., vol. 19, pp. 1193–1207, Oct. 2022.

56. J. A. Kelly, A. N. Olson, K. Neupane, S. Munshi, J. San Emeterio, L. Pollack, M. T. Woodside, and J. D. Dinman, “Structural and functional conservation of the programmed -1 ribosomal frameshift signal of SARS coronavirus 2 (SARS-CoV-2),” J. Biol. Chem., vol. 295, pp. 10741–10748, 2020.

57. J. F. Atkins, G. Loughran, P. R. Bhatt, A. E. Firth, and P. V. Baranov, “Ribosomal frameshifting and transcriptional slippage: From genetic steganography and cryptography to adventitious use,” Nucleic Acids Res., vol. 44, no. 15, pp. 7007–7078, 2016.

58. C.-F. Hsu, K.-C. Chang, Y.-L. Chen, P.-S. Hsieh, A.-I. Lee, J.-Y. Tu, Y.-T. Chen, and J.-D. Wen, “Formation of frameshift-stimulating RNA pseudoknots is facilitated by remodeling of their folding intermediates,” Nucleic Acids Res., vol. 49, pp. 6941–6957, 2021.

59. P. Z. Johnson and A. E. Simon, “RNAcanvas: interactive drawing and exploration of nucleic acid structures,” Nucleic Acids Res., vol. 51, pp. W501–W508, 2023.

