## Supplemental Information for "An intricate balancing act: Upstream and downstream frameshift co-regulatory elements"

(Dated: 21 June 2024)

[a)] Both contributed equally to this work as first co authors.

[b)] To whom correspondence should be address

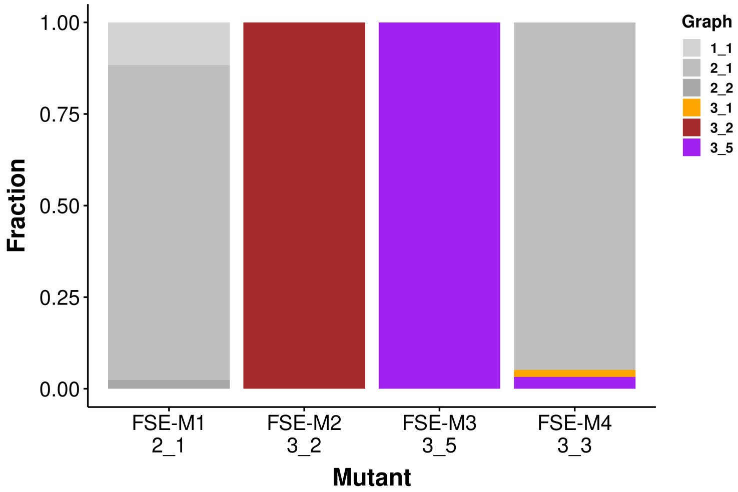


FIG. S1. Folding of 77-nt double mutants in our previous work[1] embedded in 114-nt and examined by DMS-MaP in Pekarek et al.[2]. The fractions of RNA folds are calculated from DREEM clustering (K=3) and ShapeKnots predictions weighted by Boltzmann factor. The topology of 77-nt FSE regions are represented via dual graph IDs. The target fold of each mutant is labeled under it name. Note that FSE-M1, M2 and M3 embedded in long constructs predict folds as designed for 77-nt, but FSE-M4 yield mostly 2_1 because the S3 loop that should be pseudoknotted with the downstream stem is flexible/unstable according to the DMS data.

### REFERENCES

[1]T. Schlick, Q. Zhu, S. Jain, and S. Yan, "Structure-altering mutations of the SARS-CoV-2 frameshifting RNA element," Biophys. J. **120**, 1040–1053 (2021).

[2]L. Pekarek, M. M. Zimmer, A.-S. Gribling-Burrer, S. Buck, R. Smyth, and N. Caliskan, "Cis-mediated interactions of the SARS-CoV-2 frameshift RNA alter its conformations and affect function," Nucleic Acids Res. **51**, 728–743 (2023).

[3]C. E. Hajdin, S. Bellaousov, W. Huggins, C. W. Leonard, D. H. Mathews, and K. M. Weeks, "Accurate SHAPE-directed RNA secondary structure modeling, including pseudoknots," Proc. Natl. Acad. Sci. U.S.A. **110**, 5498–5503 (2013).

[4]P. J. Tomezsko, V. D. A. Corbin, P. Gupta, H. Swaminathan, M. Glasgow, S. Persad, M. D. Edwards, L. Mcintosh, A. T. Papenfuss, A. Emery, R. Swanstrom, T. Zang, T. C. T. Lan, P. Bieniasz, D. R. Kuritzkes, A. Tsibris, and S. Rouskin, "Determination of RNA

TABLE S1. Full sequences of the mutants in this work. Mutations are colored and in bold. The wildtype, M3_3, M3_5, M3_6 and M3_5$^+$ are 118-nt; M3_6$^+$ is 132-nt; and M2_2 is 156-nt.

| System | Sequence |
|---|---|
| Wildtype | AACCCAUGCUUCAGUCAGCUGAUGCACAAUCGUUUUUAAAC<br>GGGUUUGCGGUGUAAGUGCAGCCCGUCUUACACCGUGCGGCACAGGCACUAGUACUGAUGUCGUAUACAGGGCUUUU |
| M3_3 | AACCCAUGCUUCAGUCAGCUGAUGCACAAUCGUUUUUAAAC<br>GGG**C**UUGCGGUGUAAGUGCAGCCCGUCUUACACCGUGCGGCACAGGCACUAGUACUGAUGUCGUAUACAG**AU**CUUUU |
| M3_5 | AACCCAUGCUUCAGUCAGCUGAUGCACAAUCGUUUUUAAAC<br>GGGUUUGCGGUGUAAGUGCAGCCCGUCUUACACCGUGCGGCACAGGCACUAGUACUGAUGUCGUAUACAGG**C**C**C**UUU |
| M3_6 | AACCCAUGCUUCAGUCAGCUGAUGCACAAUCGUUUUUAAAC<br>GG**UA**UUGCGGUGUAAGU**AA**AGCCCGUCUUACACCGUGCGGCACAGGCACUAGUACUGAUGUCGUAUA**AC**GGGCUUUU |
| M3_5$^+$ | AACCCAUGCUUCAGUCAGCUGAUGCACAAUCGUUUUUAAAC<br>GGGUUUGCGGUGUAA**UG**GCA**A**C**UA**GUCUUACACCGUGCGGCACAGG**A**ACUA**U**UACUGAUGUCGUAUACAGG**C**C**C**UUU |
| M3_6$^+$ | UGAUCAACUCCGCG<br>AACCCAUGCUUCAGUCAGCUGAUGCACAAUCGUUUUUAAAC<br>**UUU**UUUGCGGUGUAAGUGCAGCCCGUCUUACACCGUGCGGCACAGGCACUAGUACUGAUGUCGUAUACAGGGCUUUU |
| M2_2 | ACUCCGCGAACCCAUGCUUCAGUCAGCUGAUGCACAAUCGUUUUUAAAC<br>GGGUUUGCGGUGUA**UA**UGCAGCCCGUCUU**U**CACCGUGCGGCACAGGCACUAGUACUGAUGUCGUAUACAGGGCUUUU<br>GACAUCUACAAUGAUAAAGUAGCUGGUUUU |

TABLE S2. Secondary structures predicted by eight software packages for SARS-CoV-2 77-nt FSE and motif strengthening mutants. MFE structures are presented in dual graph notations.

| Software | 77WT | M3_6 | M3_5 | M3_3 |
|---|---|---|---|---|
| **ProbKnot** | 3_5 | ✓3_6 | ✓3_3 | ✓3_5 |
| **Pknots** | ✓3_6 | ✓3_6 | ✓3_3 | ✓3_5 |
| **Ipknot** | 2_1 | ✓3_6 | No graph match | ✓3_5 |
| **vsfold5** | 4_7 (3_3-containing) | ✓3_6 | 4_3 (3_3-containing) | 4_3 (3_3-containing) |
| **pKiss** | ✓3_6 | ✓3_6 | ✓3_3 | ✓3_5 |
| **Vfold2D** | ✓3_6 | 3_3 | ✓3_3 | ✓3_5 |
| **Knotty** | 4_7 (3_3-containing) | ✓3_6 | 4_7 (3_3-containing) | ✓3_5 |
| **NUPACK** | ✓3_6 | ✓3_6 | ✓3_3 | ✓3_5 |
| **ShapeKnots** | 3_5 | ✓3_6 | 2_1 | ✓3_5 |

structural diversity and its role in HIV-1 RNA splicing," Nature **582**, 438–442 (2020).

[5] E. Morandi, I. Manfredonia, L. M. Simon, F. Anselmi, M. J. van Hemert, S. Oliviero, and D. Incarnato, "Genome-scale deconvolution of RNA structure ensembles," Nat. Methods **18**, 249–252 (2021).

TABLE S3. Boltzmann fractions computed for 77-nt, 87-nt and 114-nt SARS-CoV-2 FSE constructs using DREEM and DRACO.

| System | Motif | DREEM | DRACO |
|---|---|---|---|
| Wildtype 77-nt | 3_6 | 0.935 | 0.657 |
| | 3_5 | 0.0006 | 0.002 |
| | 3_3 | 0.057 | 0.145 |
| | 2_1 | 0 | 0.195 |
| Wildtype 87-nt | 3_6 | 0.89 | 0.16 |
| | 3_5 | 0.03 | 0.13 |
| | 3_3 | 0.06 | 0.57 |
| | 2_1 | 0 | 0.14 |
| Wildtype 114-nt (Pekarek et al. FSE-V1) | 3_3 | 0.95 | NA |
| | 3_5 | 0.03 | NA |
| | 2_1 | 0.01 | NA |


[6] J. N. Zadeh, C. D. Steenberg, J. S. Bois, B. R. Wolfe, M. B. Pierce, A. R. Khan, R. M. Dirks, and N. A. Pierce, "NUPACK: Analysis and design of nucleic acid systems," J. Comput. Chem. **32**, 170–173 (2011).

[7] E. Rivas and S. R. Eddy, "A dynamic programming algorithm for RNA structure prediction including pseudoknots11Edited by I. Tinoco," J. Mol. Biol. **285**, 2053–2068 (1999).

[8] S. Janssen and R. Giegerich, "The RNA shapes studio," Bioinformatics **31**, 423–425 (2015).

[9] T. C. T. Lan, M. F. Allan, L. E. Malsick, J. Z. Woo, C. Zhu, F. Zhang, S. Khandwala, S. S. Y. Nyeo, Y. Sun, J. U. Guo, M. Bathe, A. Näär, A. Griffiths, and S. Rouskin, "Secondary structural ensembles of the SARS-CoV-2 RNA genome in infected cells," Nat. Commun. **13**, 1128 (2022).

[10] N. C. Huston, H. Wan, M. S. Strine, R. de Cesaris Araujo Tavares, C. B. Wilen, and A. M. Pyle, "Comprehensive in vivo secondary structure of the SARS-CoV-2 genome reveals novel regulatory motifs and mechanisms," Mol. Cell **81**, 584–598.e5 (2021).

[11] C. Cao, Z. Cai, X. Xiao, J. Rao, J. Chen, N. Hu, M. Yang, X. Xing, Y. Wang, M. Li, B. Zhou, X. Wang, J. Wang, and Y. Xue, "The architecture of the SARS-CoV-2 RNA genome inside virion," Nat. Commun. **12**, 3917 (2021).

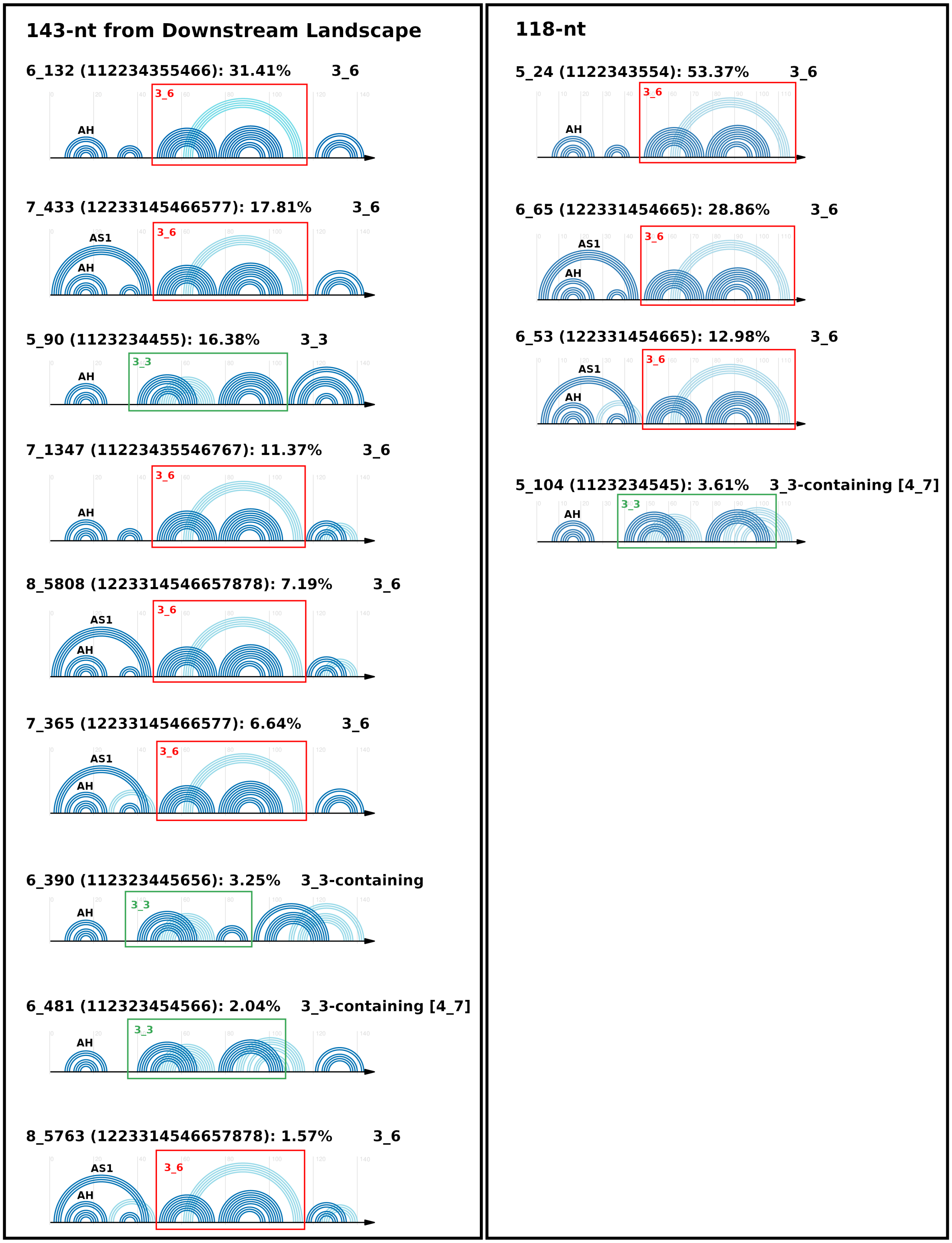


FIG. S2. Secondary structures of 143-nt and 118-nt FSE from the downstream landscape. Corresponding fractions are labeled for each motif. The dominant motifs are AH + 3_6 for 95.92% at 118-nt and for 75.99% at 143-nt. The minor motifs are AH + 3_3 for 3.61% at 118-nt and for 21.67% at 143-nt.

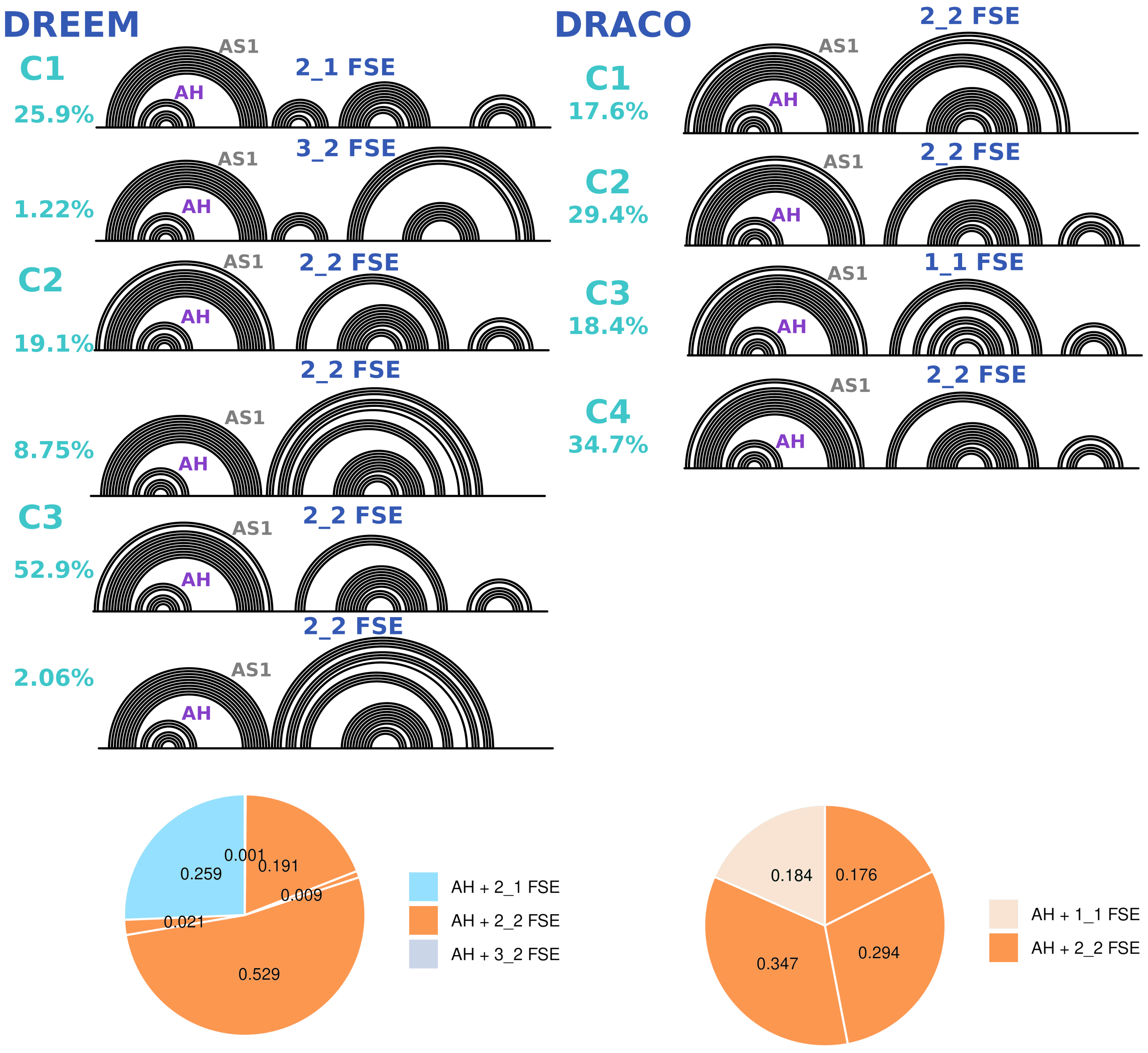


FIG. S3. The 156-nt construct is investigated by DMS-MaPseq and the folds are predicted by ShapeKnots[3] guided by DMS reactivity profiles. DREEM[4] and DRACO[5] have been applied to obtain clusters of alternative folds and produce DMS reactivity profile. DREEM were run with the setting K(number of clusters)=3, and DRACO produces 4 clusters.

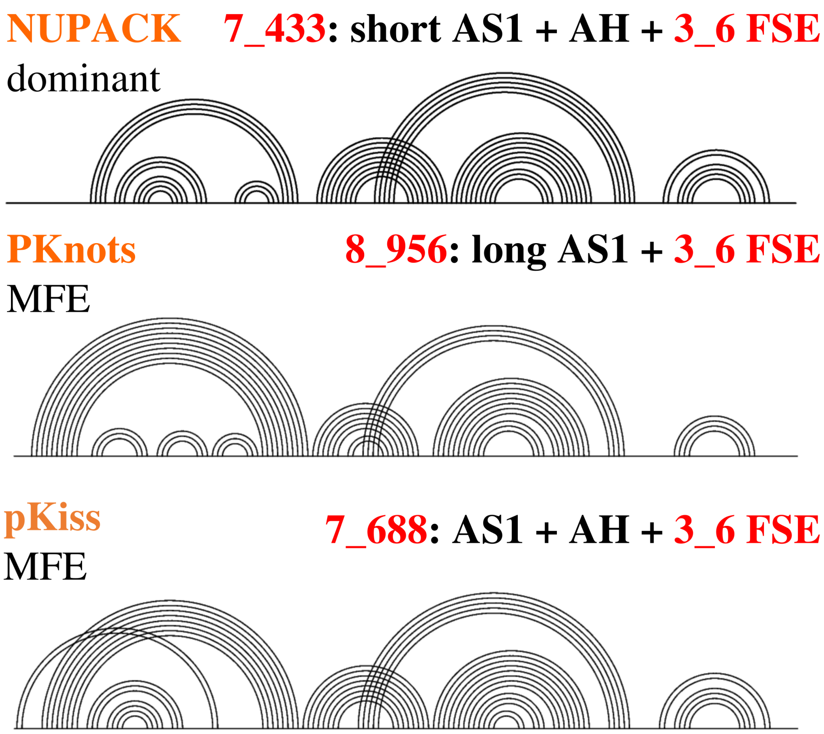


FIG. S4. Predictions of 156-nt FSE construct by selected software packages (NUPACK[6], PKNOTS[7] and pKiss[8]).

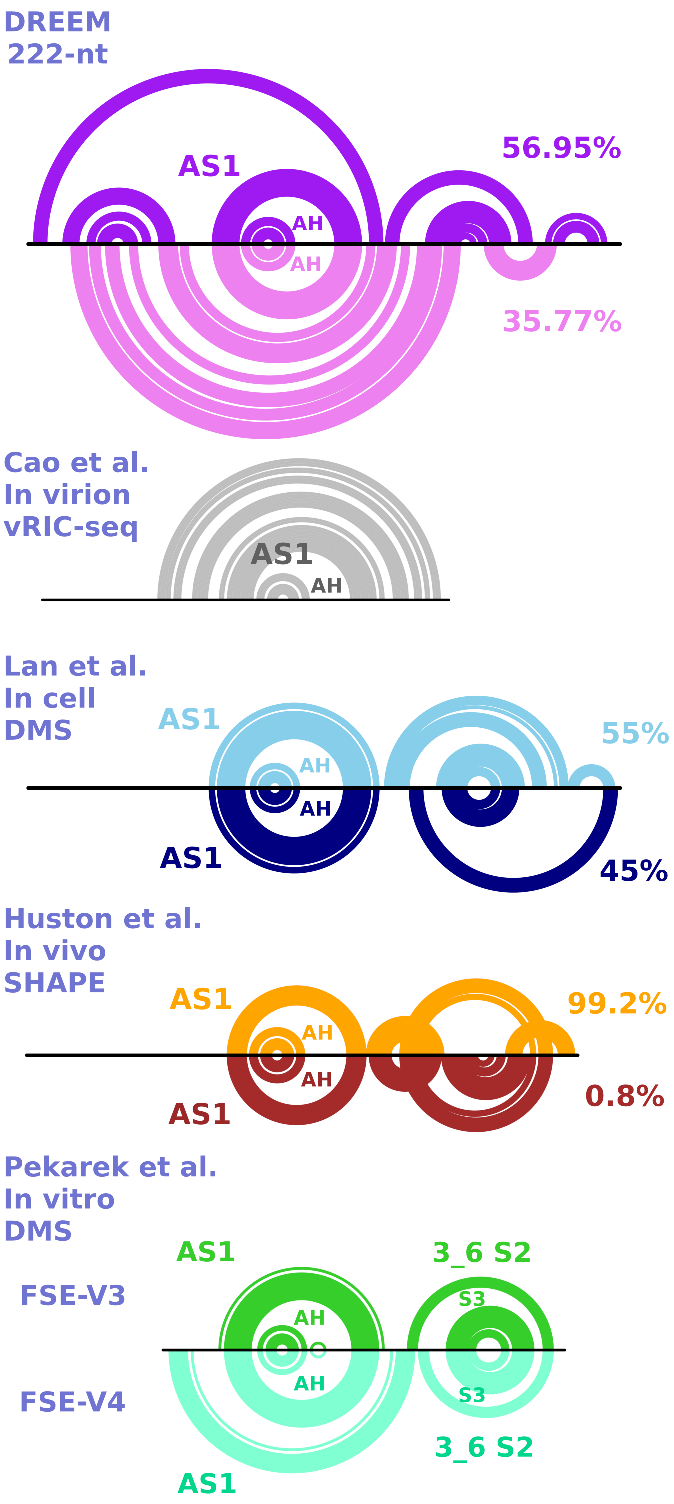


FIG. S5. 222-nt FSE-containing construct characterized by DMS and structures characterized in different experimental conditions in the literature[9–11].

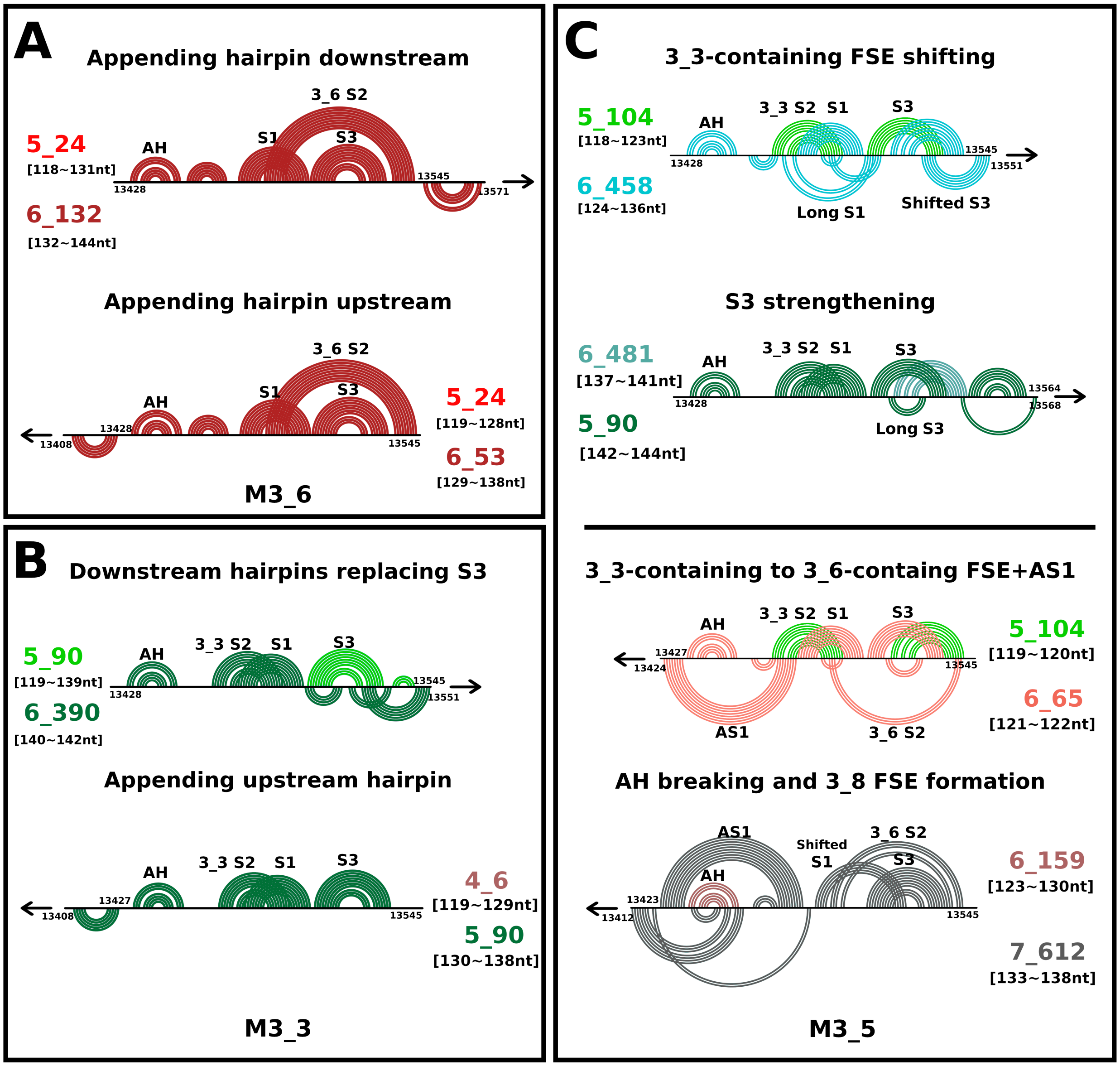


FIG. S6. Transitions of 2D folding from the dominant motifs of FSE mutant systems in this work with additional downstream and upstream sequence are shown in arc plots, for (A-C) the motif strengthening mutants M3_6 (A), M3_3 (B), and M3_5 (C). The 2D arc plots for two motifs in different colors are overlapped to show the differences in RNA folding. 3_6 FSE containing motifs are colored in red; similarly, 3_3 FSE containing motifs are in green, and 3_8-containing motifs are in gray. For the M3_6 (A) and M3_3 (B), two transitions are displayed, top for downstream expansion and bottom for upstream expansion (expansion indicated by the arrows). For M3_5 (C), four transitions are displayed, with the top two transitions indicating downstream sequence expansion and the bottom two indicating upstream sequence expansion. The overlapped arc plots depict each transition, with added pairs in the second motif at the bottom and common stems at the top.

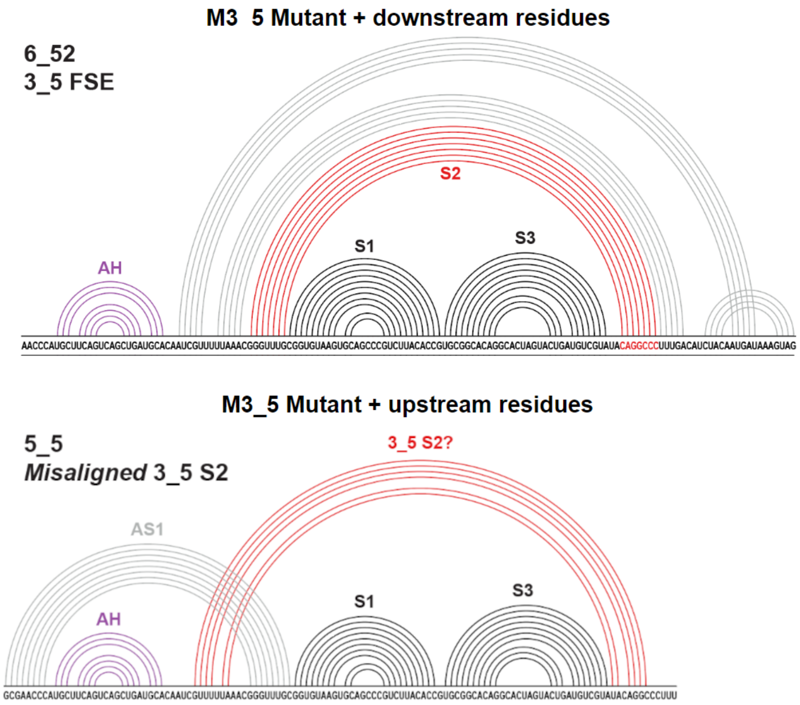


FIG. S7. Motifs containing AH and 3_5 FSE that rarely appear for M3_5 system.
